# Identification of an isoflavonoid transporter required for the nodule establishment of the *Rhizobium*-*Fabaceae* symbiotic interaction

**DOI:** 10.1101/2021.07.30.452494

**Authors:** Wanda Biała-Leonhard, Laura Zanin, Stefano Gottardi, Rita de Brito Francisco, Silvia Venuti, Fabio Valentinuzzi, Tanja Mimmo, Stefano Cesco, Barbara Bassin, Enrico Martinoia, Roberto Pinton, Michał Jasiński, Nicola Tomasi

## Abstract

Nitrogen (N) as well as Phosphorus (P) are key nutrients determining crop productivity. Legumes have developed strategies to overcome nutrient limitation by e.g., forming a symbiotic relationship with N-fixing *rhizobia* and the release of P-mobilizing exudates and are thus able to grow without supply of N or P fertilizers. The legume-rhizobial symbiosis starts with root release of isoflavonoids, that act as signaling molecules perceived by compatible bacteria. Subsequently, bacteria release nod factors, which induce signaling cascades allowing the formation of functional N-fixing nodules.

We report here the identification and functional characterization of a plasma membrane-localized MATE-type transporter (LaMATE2) involved in the release of genistein from white lupin roots.

The *LaMATE2* expression in the root is upregulated under N deficiency as well as low phosphate availability, two nutritional deficiencies that induce the release of this isoflavonoid. *LaMATE2* silencing reduced genistein efflux and even more the formation of symbiotic nodules, supporting the crucial role of LaMATE2 in isoflavonoid release and nodulation. Furthermore, silencing of LaMATE2 limited the P-solubilization activity of lupin root exudates. Transport assays in yeast vesicles demonstrated that LaMATE2 acts as a proton-driven isoflavonoid transporter.

## INTRODUCTION

One of the major challenges of sustainable agriculture comprises the production of high-quality plant material with preservation of soil components and reduced application of chemical fertilizers without penalizing yield. Nitrogen (N) and phosphorus (P) are limiting nutrients in most natural soils (Tilman, 1999). High input of N-fertilizers is required to sustain crop growth in conventional agriculture. However, this feature may contaminate soils and groundwater and markedly contribute to the release of greenhouse gases (Tilman et al., 2001). Much of the P in soils is not available to plants due to its tendency to interact with calcium and magnesium salts or iron and aluminum oxides. It is mainly present as sparingly soluble rock phosphate, or it is immobilized in slowly mineralizable P-containing organic compounds, such as phytates. Phosphorous is a non-renewable resource which is mined at an increasing rate to meet the demand for fertilizers (Vance et al., 2003).

*Leguminous* plants (*Fabaceae*) such as soybean and lupin, have evolved several strategies to survive in low nutrient soils. In the case of P, in many ecosystems and, in particular, in acidic soils, the plant’s response consists mainly in the association with mycorrhizal fungi or the formation of particular root structures, such as cluster roots (Purnell, 1960; Neumann and Martinoia, 2002; Lambers et al., 2015). Cluster roots release huge amounts of exudates into the rhizosphere which are mainly composed of carboxylates and flavonoids. Flavonoids are involved both in the mobilization of nutrients and in the modulation of soil microbial activities (Cesco et al., 2010). The production and release of flavonoids, are also essential for the establishment of the symbiotic interaction between legumes and N-fixing bacteria such as *Ensifer*, *Bradyrhizobium* or *Mesorhizobium*, leading to the fixation of atmospheric N. Moreover it has been hypothesized that flavonoids are involved in the initiation of the nodule through their action on the plant hormone auxin and could thus play a developmental role in addition to their action as nod gene regulators (Subramanian et al., 2006; Wasson et al., 2006; Li et al., 2016).

The mutualistic fungal and bacterial symbionts are striking examples of soil microorganisms that have successfully coevolved with their hosts since plants adapted to terrestrial ecosystems. They promote plant growth by facilitating the acquisition of scarce nutrients. The most commonly established symbiosis in plants is the mycorrhizal association, with 80–90% of all land-plant species able to enter this interaction. Around 100 Mio years ago, certain angiosperms evolved a bias toward the evolution of nodulation with the so-called N-fixing soil bacteria. Among those angiosperms are legumes (*Fabales*) and one non-legume genus, *Parasponia* (*Cannabaceae*, *Rosales*) which can establish mutualistic symbioses with *Rhizobia*, a polyphyletic group of proteobacteria and diverse group of plants belonging to the orders *Fagales*, *Rosales*, and *Cucurbitales* which can associate symbiotically with filamentous actinobacteria of the genus *Frankia* (Martin et al., 2017). By forming symbiotic associations, plants obtain mineral nutrients. In turn, they supply the symbiont with organic compounds, sugars and lipids in the case of mycorrhiza, mostly carboxylates to N-fixing bacteria (Udvardi and Poole, 2013; Jiang et al., 2017; Luginbuehl et al., 2017). The establishment of the symbiosis is a complex event and requires coordinated regulation of the corresponding genes and release of signaling molecules into the rhizosphere. For the legume-*rhizobia* symbiosis, it is expected that a flavonoid transporter must be present in the plasma membrane of root cells to release isoflavonoids into the rhizosphere (Sugiyama et al., 2007). Up to now, transporters for flavonoids have been mainly described at the vacuolar membrane (Zhao, 2015). Furthermore, an ABC (ATP-Binding Cassette) transporter from *Medicago* was shown to transport flavonoids. However, this transporter is localized in the vasculature (Biala et al., 2017). Using a biochemical approach Sugiyama and coworkers (2007) presented evidence that, in soybean, genistein transmembrane transport is mediated by an ABC-type transport system. But its contribution to genistein root release and the legume-*rhizobia* symbiosis establishment is still unclear.

Despite all the research performed on this symbiotic interaction, a transporter releasing flavonoids into the rhizosphere and initiating the first step of this symbiosis awaits its identification.

## MATERIALS AND METHODS

### Plant growth and transformation

White lupin seeds (*Lupinus albus* L. cv. Amiga; Südwestdeutsche Saatzucht, Rastatt, Germany) were soaked for 24 hours in aerated water and germinated on a plastic net placed at the surface of an aerated 0.5 mM CaSO_4_ solution in a growth chamber at 25 °C in the dark. Thereafter, 7-day-old seedlings were transferred to a hydroponic system, containing a P-free nutrient solution (μM): 5000 Ca(NO_3_)_2_, 1250 MgSO_4_, 1750 K_2_SO_4_, 250 KCl, 20 Fe(III)EDTA, 25 H_3_BO_4_, 1.25 MnSO_4_, 1.5 ZnSO_4_, 0.5 CuSO_4_, 0.025 (NH_4_)_6_Mo_7_O_4_. Phosphorus-deficient plants were grown on P-free nutrient solution, while 0.25 mM KH_2_PO_4_ were added to nutrient solution for P-sufficient condition. Plants were grown under controlled conditions for 4 weeks (day/night photoperiod, 16/8 h; radiation, 220 µE m^−2^s^−1^; day/night temperature, 25/20 °C; relative humidity, 70-80%).

In P-deficiency, lupin plants modify the root architecture developing particularly root structures, called cluster roots or proteoid roots. In order to differentiate the developmental stages of root clusters (Figure S1), the root system was immerged in a pH-indicator solution (0.04% w/v bromocresol purple). Depending on their morphology and capability to acidify the solution, at the end of the growing period different regions of cluster root were sampled from P-deficient plants (juvenile, immature, mature, senescent), as described by Massonneau and coworkers (2001). Root apices and cluster root parts were sampled from a pool of 12-16 P-sufficient and P-deficient plants. The samples were immediately frozen in liquid nitrogen for the RNA extraction or were rinsed twice in 0.5 mM CaSO_4_ solution and the root exudates were collected for one hour in 0.5 mM CaSO_4_ 10 mM 2-[N-Morpholino]ethanesulfonic acid (MES)-KOH pH 6.0 at a ratio 1:10 W/V. The samples were conserved at –80 °C until processing. Three independent experiments were performed.

To investigate the plant response to N-deficiency, white lupin seeds were grown for two weeks on watered Whatman paper in Petri dishes and then transferred to the magenta boxes containing PFR N-free solution described by Stróżycki and coworkers (2003). For N-sufficient conditions KNO_3_ (0.1 mM), NH_4_H_2_PO_4_ (5 mM), Ca(NO_3_)_2_ (2.46 mM) were added to the PFR medium. Plants were grown for further two weeks under controlled conditions (day/night photoperiod, 16/8 h; radiation, 220 µE m^−2^s^−1^; day/night temperature, 23/20 °C; relative humidity, 70-80 %). The samples were collected at 7 and 14 days after the transfer in N-deficient condition. Two biological repetition and 4/8 technical repetition were performed.

White lupin seedlings were transformed using *Agrobacterium rhizogenes* ARqua1 strain (Quandt et al., 1993) carrying binary vector pRedRoot::*LaMATE2 RNAi* or empty vector (EV) pRedRoot. After germination 5 mm root tips were removed from radicles. The sectioned surface was coated with *A. rhizogenes* and seedlings were placed on solid Fahraeus medium supplemented with kanamycin (15 mg L^−1^).

Nitrogen-deficient plants were grown and analyzed as described above. For P-deficiency, after 3 weeks post germination, plants were transferred to P-free nutrient solution (as described above). After 2 weeks, white lupin plants were moved in hydroponic solutions in P-free nutrient solution and grown for 3 additional weeks. *LaMATE2* expression analyses were performed, and root release was collected as described previously on the fully developed cluster root (immature, mature). Six biological replicates were performed for each sample.

### Nodulation and effects of genistein exogenous addition

Three-week-old lupin composite plants were transferred to the pots filled with sterile perlite (0.75 L). Plants were nourished with PFR N-free solution. After 2 week of N-deficiency plants were inoculated with *B. japonicum* (strain UPP 133, (Stępkowski et al., 2011)) and grown for further two weeks under controlled conditions (day/night photoperiod, 16/8 h; radiation, 220 µE m^−2^s^−1^; day/night temperature, 23/20 °C; relative humidity, 70-80 %). For complementation experiments 1 h before inoculation with *B. japonicum* plants were additionally supplemented with 1 µM genistein solution. Fourteen-day post-inoculation plants were removed from pots, nodules were counted for each single root, and roots were collected for each plant. Afterwards all roots of single plant were grounded and divided into aliquots dedicated for gene expression and metabolomic analysis. Only plants revealing expression of marker gene encoding fluorescent protein DsRed were considered during analysis. These experiments were performed 6, 8 or 21.

### Mobilization of P from vivianite

The vivianite suspension (Fe_3_(PO_4_)_2_.8H_2_O) was prepared by mixing H_3_PO_4_ with FeSO_4_·as described by Eynard et al. (1992). A solution of 0.035 M H_3_^32^PO_4_ and 0.05 M FeSO_4_·7H_2_O were brought to pH 6.0 with 0.05 M KOH under stirring. A bluish suspension of vivianite was obtained, after precipitation via centrifugation (8000 g, 10 min), the precipitate was washed (by 3 successive centrifugations and decantations) with deionized water. Mobilization of PO_4_^2-^ from Vivianite (Fe_3_(PO_4_)_2_.8H_2_O) by exudates after purification with the SPE methods, in order to remove carboxylates, from *LaMATE2 RNAi* or empty vector (EV) pRedRoot transformed P-deficient roots was performed as followed: one mL of a suspension containing ^32^P-vivianite (70 μmol PO_4_, specific activity 60 KBq μmol^−1^ P) was transferred into a dialysis tube (ZelluTrans/Roth 6.0, ∅ 16mm, exclusion limit of 4÷6 kDa, ROTH) and mixed with 5 mL of mobilization solution [0.5mM CaSO_4_, 20mM Mes-KOH (pH 6.0), 50 µl methanol]. Thereafter, the dialysis tube was transferred into continuously mixed 34 ml of mobilization solution. At the beginning of the experiment, exudates corresponding to the amount release for 1 hour by 1 gram of juvenile cluster root tissues (Figure 1) were added to the external solution. After 15, 30, 45 and 60 min, samples from the solution outside the dialysis tube were collected and the amount of ^32^P was measured by liquid scintillation counting. The ^32^P mobilization was estimated from the difference between the ^32^P concentration measured in the presence of the mock and in the absence of exudates and was expressed as µmol P h^−1^.

**Figure 1.**
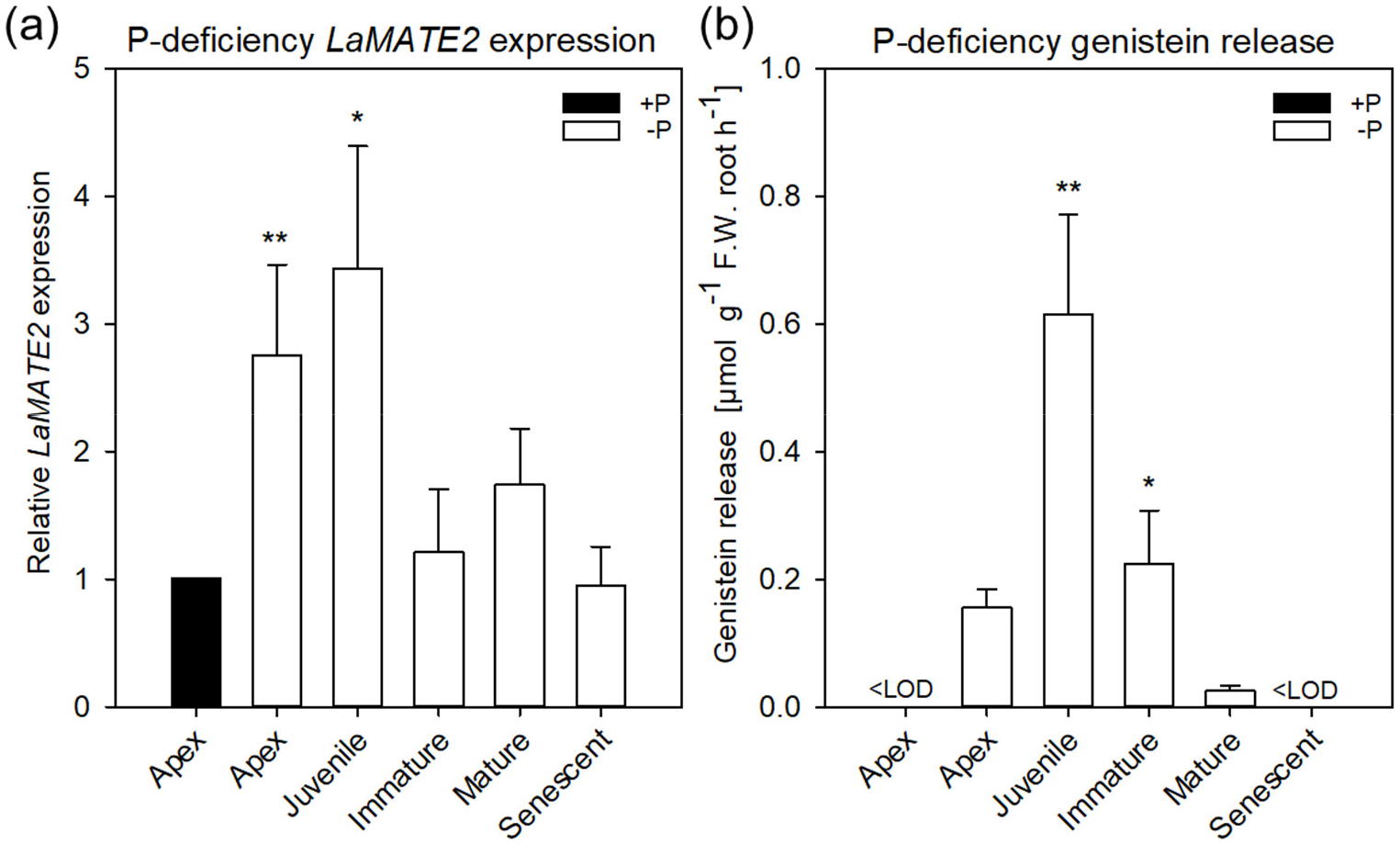
*LaMATE2* expression level and genistein release in roots of P-deficient white lupin. *LaMATE2* expression analyses in white lupin grown under control (+N+P, black bars) condition or P deficiency (+N-P, white bars) (**a**). Gene expression was evaluated in control apex (+N+P), P-deficient apex and cluster roots (separated depending on developing stages: Juvenile, Immature, Mature, Senescent cluster-root stages; example of cluster root shown in Figure S1). Expression levels are shown relative to the *LaMATE2* expression level of the root apex under control conditions (+N+P). Release of genistein from different root tissues of 4-week-old P-deficient plants, release from the control apex was below detectable value <LOD (**b**). Data are means +SD (* refer to statistically significant differences among the mean value of the sample *vs* control or -P apex for genistein release, ANOVA Holm–Sidak, N=6, *P <0.05, ** P <0.01).

### Extraction and LC/MS analysis

Frozen root tissue (500 mg) was grounded and extracted with 80% methanol. Root exudates were extracted from the medium by SPE (Solid Phase Extraction) method using octadecylosilane matrix and methanol according to Staszków and coworkers (2011). Luteolin was used as an internal standard. Samples were analyzed by liquid chromatography-electrospray ionization tandem mass spectrometry (LC/ESI/MS) using a Waters UPLC coupled with Bruker micrOTOF-Q mass spectrometer. The analysis was performed in a gradient mobile phase consisting of 0.5% formic acid (v/v) in water (A) and 0.5% formic acid (v/v) in acetonitrile (B). The m/z range of the recorded spectra was 50-1000. Analyses were performed in the ion-positive mode.

### RNA extraction and cDNA synthesis

RNA extractions were performed using the InviTrap Spin Plant RNA Mini Kit (Stratec Molecular, Berlin, Germany) following manufacturer’s instructions and contaminant genomic DNA was removed using 10 U of DNase I (GE Healthcare, Munich, Germany). The quantity and the quality of RNA was checked using a spectrophotometer, followed by a migration in a 1 % agarose gel. One microgram of total RNA for each sample was retro-transcribed using 1 pmol Oligo d(T)23 (Sigma Aldrich, Saint Louis, USA) and 10 U M-MulV RNase H (Finnzymes, Helsinki, Finland) following manufacturers’ instruction.

### Isolation of the *LaMATE2* sequence

The partial sequence was isolated via a cDNA-AFLP approach starting from RNA extracted from juvenile cluster roots which were compared to mature and senescent cluster roots, for details see Massonneau and coworkers (2001). The full Open Reading Frame (ORF) of *LaMATE2* (*LaMATE2*_ORF_) was isolated from the cDNA of juvenile cluster root tissues from P-deficient plants using the 5’/3’ RACE Kit (2nd generation, Roche Diagnostics S.p.a., Monza, Italy) following the manufacturer’s instructions. *LaMATE2*_ORF_ sequence was cloned in pGEM-T easy vector (for primers see Table S1, Promega Italia Srl, Milan, Italy) and deposited in the NCBI database (KY464927).

### Gene expression analysis

The reaction was performed by adding 0.1 µl of cDNA to RT complete reaction mix, Fluocycle^TM^ sybr green (20-µl final volume; Euroclone, Pero, Italy). Specific primers were designed for the target, the two closest homologues (www.whitelupin.fr; Hufnagel *et al*., 2020) and the housekeeping gene using Primer3 software (Koressaar and Remm, 2007; Untergasser et al., 2012) and were synthesized by Sigma Aldrich (Table S1). Gene expression analyses were performed using CFX96^TM^ Real-Time System (C1000TM Thermal Cycler, BioRad) and CFX Manager™ Software (v 2.0, BioRad). The two closest homologues (*Lalb_Chr06g0165721, Lalb_Chr02g0145611*) have a low expression in the root tissues and their expression levels is shown in Figure S5. Data were normalized in respect to the transcript level of the housekeeping gene (*Ubiquitin* gene, *LaUBI*) using the 2^−ΔΔCT^ method (Livak and Schmittgen, 2001). The efficiency of amplification was calculated using R program (version 2.9.0; http://www.r-project.org/) with the qPCR package (version 1.1-8), following the authors’ indications (Ritz and Spiess, 2008). Six biological replicates were performed for each sample.

### Genistein transport by LaMATE2 in *Saccharomyces cerevisiae* vesicles

The construction of the yeast expression vector pNEV (Sauer and Stolz, 1994) containing *LaMATE2*_ORF_ was performed by amplifying *LaMATE2*_ORF_ sequence with primers having both a *NotI*-restriction sites at the 5’end. The cloning of the PCR product (*LaMATE2*_ORF_) was performed into the *NotI* site of pNEV (pNEV::*LaMATE2*_ORF_), the orientation was verified by sequencing. The transformation of competent yeast cells (*S. cerevisiae* YPH499 strain) was performed following a standard procedure (Gietz and Woods, 2002) and transformants were selected on synthetic dextrose minimal medium lacking uracil (SD-Ura medium; Burke *et al*., 2000).

The yeast cells, strain YPH499, were transformed with the pNEV empty vector or pNEV::LaMATE2_ORF_ and selected on liquid-SD Ura-medium. Thereafter, cells were incubated in YPD medium for 30 min, collected by centrifugation, and digested with lyticase (1,000 U g^−1^ fresh weight cells; Sigma Aldrich), and subsequently microsomal vesicles were isolated as described by Klein and coworkers (2002). Transport assays were performed to study the genistein transport using the rapid filtration technique with nitrocellulose filters (0.45 µm pore size; Millipore, Millipore Co., Bedford, USA). The transport experiment was carried out in presence of isolated vesicles, transport buffer (0.4 M glycerol, 0.1 M KCl, 1 mM DTT, 1 mM EDTA, 5 mM ATP, 10 mM MgCl_2_, 10 mM creatine phosphate, 0.1 mg ml^−1^ creatine kinase, 20 mM Tris-MES pH 7.4) and 5 µM of labelled ^3^H-genistein (American Radiolabeled Chemicals, Saint Louis, USA; 1850 Bq filter^−1^). Only for the kinetic experiments, the ^3^H-genistein concentration ranged from 5 µM up to 100 µM (5, 7, 10, 15, 33, 50 and 100 µM ^3^H-genistein) and the incubation time was 30 s. The kinetic parameters of ^3^H-genistein uptake were calculated by subtracting uptake rates recorded in the empty-vector vesicles. The kinetic parameters of ^3^H-genistein uptake were calculated between 7-100 µM by subtracting uptake rates recorded in the empty-vector vesicles using the Hanes– Woolf plot.

To test the dependence of genistein transport on a proton gradient, 25 mM NH_4_Cl was included in the transport buffer assay and the incubation time ranging between 15 and 120 s (15, 30, 60 and 120 s). Moreover, the capability of LaMATE2 to mediate the uptake of different flavonoids was tested by UPLC under the same experimental conditions reported above (5 µM of flavonoid: genistein, genistin, hydroxygenistein, biochanin A, daidzein, or kaempferol; the incubation time was 30 s). The mixture was loaded on a pre-wetted filter and removed by suction at the end of incubation time. The membranes were rapidly washed twice with 2 ml of ice-cold transport buffer.

The radioactive measurements were determined with a beta-counter (Tri-Carb 1900CA, Packard, Downers Grove, USA). As standards, solutions with known amounts of ^3^H-genistein were used. Vesicle protein content was quantified with BioRad Protein Assay Dye Reagent (BioRad, Hercules, CA, USA), and the data are shown as pmol ^3^H-genistein µg^−1^ protein. Data are shown as net pmol ^3^H-genistein µg^−1^ protein after removing background, i.e. unspecific adsorption of genistein onto empty-vector yeast membrane. Three independent transformations were performed for each sample.

### LaMATE2 subcellular localization in *Arabidopsis thaliana* protoplasts

For transient expression of *LaMATE2*_ORF_ in Arabidopsis protoplasts, the plasmid harboring the sequence for the *Green Fluorescent Protein* (*GFP*) was fused at the C-terminus of *LaMATE2*_ORF_ inside the pUC18-Sp-GFP6 vector (Komarova et al., 2008) using *NheI* and *SphI* restriction sites via PCR amplification. A plasmid harboring the sequence for *mCherry-fluorescent protein* was fused with *AtPIP2a*, a gene coding for an aquaporin used as a plasma membrane marker (Nelson et al., 2007). Arabidopsis protoplasts were co-transformed with both constructs, *LaMATE2*_ORF_-*GFP* and *AtPIP2a-mCherry*, using the polyethylene glycol method (Jin et al., 2001). Protoplasts were examined with a TCS SP5 confocal microscope (Leica Microsystems, Wetzlar, Germany), excited with an argon laser at 458 nm for GFP and 540–552 nm for mCherry (for GFP: excitation BP458, beamsplitter FT500, emission BP 492-511 nm; for mCherry: excitation BP 540–552, beamsplitter FT560, emission BP 575–640).

### RNAi-based silencing of *LaMATE2* gene in white lupin roots

*LaMATE2* silencing was adapted from Uhde-Stone and coworkers (2005) using binary transformation vectors, pRNAi and pRedRoot (Limpens et al., 2004). The target region (350 bp long, covering the 5’ end of *LaMATE2*_ORF_) was first amplified with PCR, and subsequently cloned into pRNAi vector between the restriction sites *NcoI*–*SwaI* and *BamHI*–*SpeI*. The cloned sequence (*LaMATE2 RNAi*) was regulated by a double CaMV35s promoter and OCS-3’ terminator. Using the *KpnI*–*PacI* restriction sites, the *LaMATE2 RNAi* cassette from the previously produced in pRNAi vector was transfered into the pRedRoot binary vector (pRedRoot::*LaMATE2 RNAi*).

### Phylogenetic and Statistical Analyses

Phylogenetic analyses were conducted using MEGA software, version 6 (Tamura et al., 2013). The tree was constructed by aligning the protein sequences by Clustal-W and the evolutionary history was inferred using the Neighbor-Joining method. The percentage of replicate trees in which the associated taxa clustered together in the bootstrap test (1000 replicates) are shown in Fig. S1 next to the branches. The tree is drawn to scale, with branch lengths in the same units as those of the evolutionary distances used to infer the phylogenetic tree. The evolutionary distances were computed using the Poisson correction method and are in the units of the number of amino acid substitutions per site.

The topology prediction of LaMATE2 was performed by using the PROTTER program (Omasits et al., 2014), while the alignment of LaMATE2 protein sequence with orthologous MATE-protein sequences was generated by Clustal-WS using Jalview software version 2 (Waterhouse et al., 2009).

Statistical significance was determined by one-way analysis of variances (ANOVA) using Holm–Sidak test, P < 0.05 or 0.01. Statistical analyses were calculated using SigmaPlot (Systat Software Inc., San Jose, CA, USA).

## RESULTS AND DISCUSSION

### Release of flavonoids and expression of *LaMATE2*

Former work from our laboratories showed that flavonoids are released mainly from the so-called juvenile, and immature (young states) cluster roots and that genistein and its derivative exudation was induced by phosphate (P) deficiency (Weisskopf et al., 2006). Furthermore, in 2001, we published research on genes differentially expressed amongst different stages of cluster roots in P-deficient white lupin (Massonneau et al., 2001). Within these genes, we identified two LaMATE-type transporters, *LaMATE1* was predominantly expressed in mature (Uhde-Stone et al., 2005), citrate excreting cluster roots, while the second (*LaMATE2*) was predominantly present in juvenile cluster roots.

To identify a flavonoid exporter possibly playing an important role in the establishment of nodules in *Fabaceae* plants, we looked first whether a link between the expression of *LaMATE2* and release of flavonoids exists. Under P-deficiency, the *LaMATE2* was predominantly expressed in the root apex and in the juvenile stage of cluster roots (Figure 1a). Nitrogen (N) deficiency, on the other hand, induced *LaMATE2* expression mainly in apex and nodule (Figure 2a). Analyses of the root exudates of white lupin revealed that genistein release was strongly induced by P deficiency, particularly in young cluster root tissues (Figure 1b), confirming previous results (Weisskopf et al., 2006). In order to see whether a similar behavior could be observed under N deficiency, plants were grown under N-deficient condition. In this case, cluster root formation was not induced. Hence, we compared the whole roots of plants grown in the presence or absence of N. Nevertheless, we could observe that under N deficiency, genistein release was also increased significantly after 1 and even more pronounced after 2 weeks (Figure 2b).

**Figure 2.**
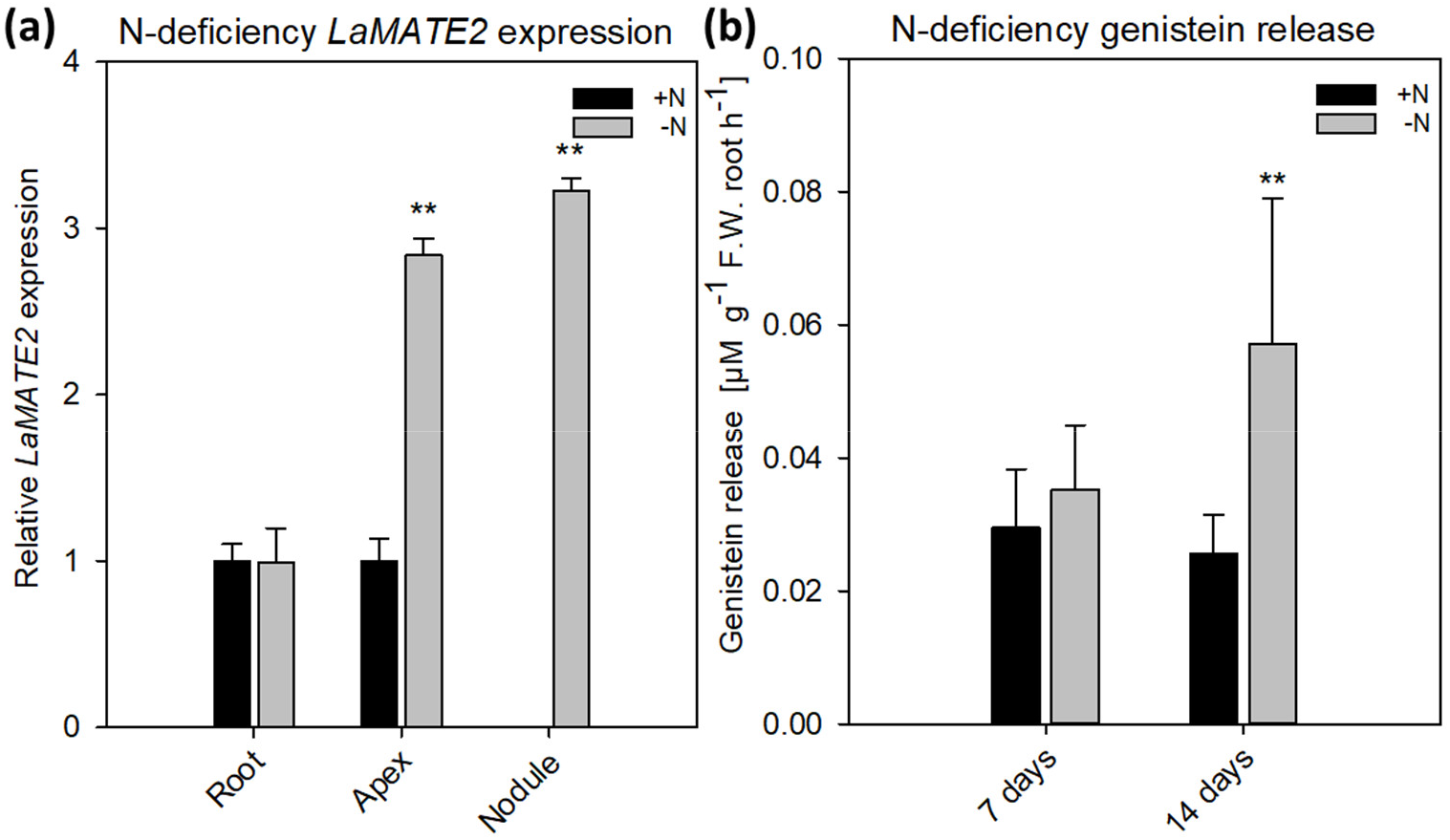
*LaMATE2* expression analyses and genistein release in roots of N-deficient white lupin. *LaMATE2* expression analyses in white lupin grown under control (+N+P, black bars) condition or N-deficiency (-N+P, dark grey bars) (**a**). Gene expression was evaluated in control and N-deficient (other) root, root apex and nodules. Expression level is shown relative to *LaMATE2* expression in control (+N+P) root. Root release of genistein from white lupin plants grown under 1- or 2-week-old N-sufficient and -deficient condition (**b**). Genistein release in 7 and 14 days of N deficiency (-N+P) or sufficiency (+N+P). Data are means +SD (* refers to statistically significant differences among the mean value of the sample *vs* control, ANOVA Holm– Sidak, N=6, * P <0.05, ** P <0.01).

### Characterization of LaMATE2

As already shown for other MATE transporters, LaMATE2 contains 12-transmembrane helical domains (Figure S2) (He et al., 2010). Indeed, LaMATE2 exhibits a good homology to other plant MATE transporters (Figure 3, S2), including the functionally characterized vacuolar flavonoid transporters MtMATE1, MtMATE2 and AtTT12 (Marinova et al., 2007; Zhao and Dixon, 2010; Zhao et al., 2011). Up to date, only few MATE transporters have been characterized in roots and most of them mediate either the efflux of citrate, such as AtFRD3, HvAACT1, SbMATE1 (Durrett et al., 2007; Furukawa et al., 2007; Magalhaes et al., 2007) or act as vacuolar flavonoid transporters in *Arabidopsis*, *Medicago* or grapevine (for a review see Zhao and Dixon (2010) and Zhao (2015)). Interestingly all citrate transporters contain a large cytosolic loop. In white lupin roots and in agreement with our previous results (Massonneau et al., 2001), the expression of another MATE transporter, LaMATE1, a homolog of the citrate transporter AtFRD3, is highest at the mature stage where a burst of citrate exudation occurs (Uhde-Stone et al., 2005; Wang et al., 2014). Also, this transporter contains a large cytosolic loop.

**Figure 3.**
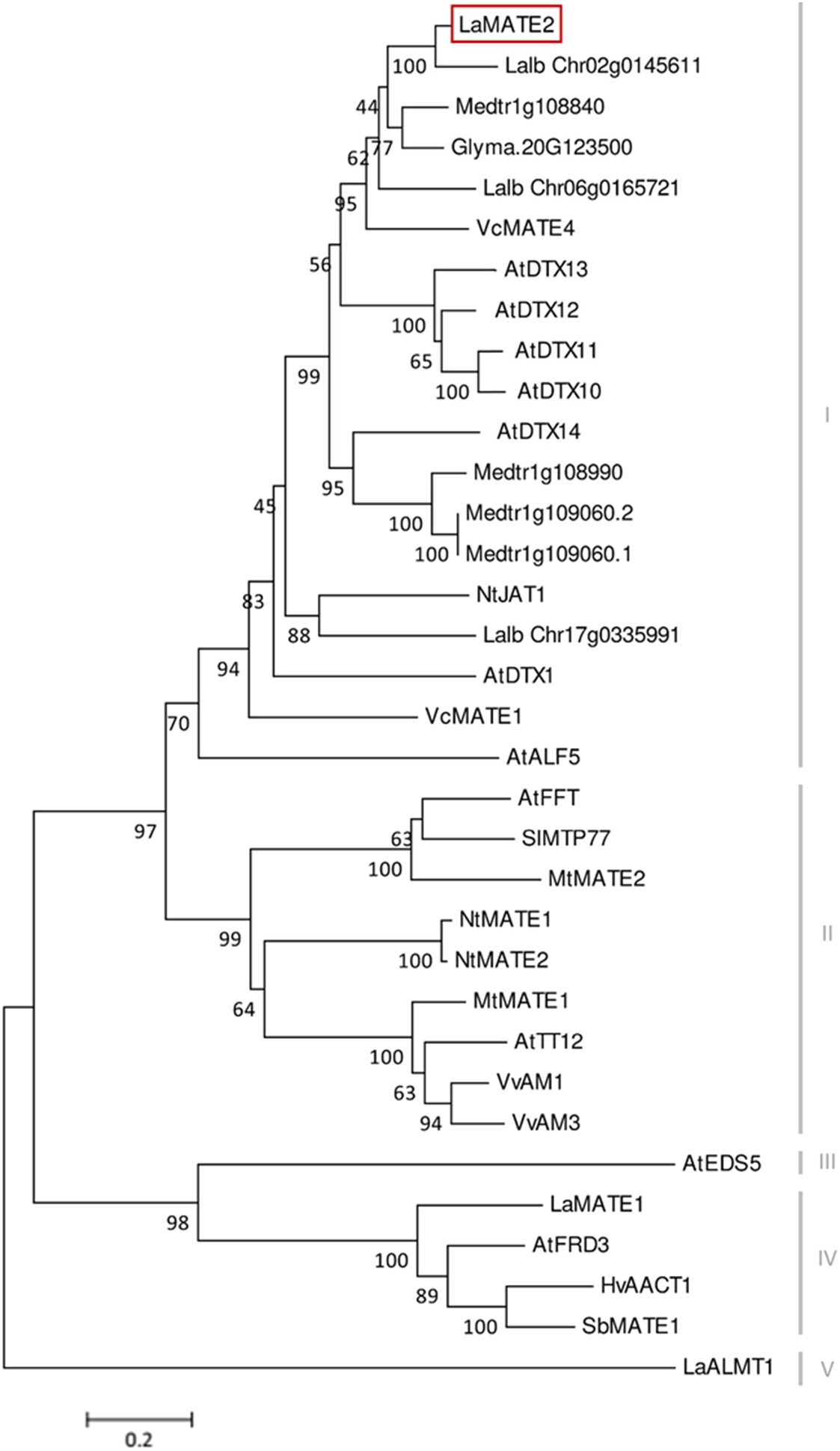
Phylogenetic tree of MATE transporters. A phylogenetic analysis was performed using the LaMATE2 amino acid sequences of *Lupinus albus* (LaMATE1, LaMATE2, Lalb_Chr17g0335991, Lalb_Chr06g0165721, Lalb_Chr02g0145611, LaALMT1); AtFRD3 AtEDS5, AtTT12, AtDTX1,10,11,12,13,14, AtALF5 and AtFFT of *Arabidopsis thaliana*; Glyma.10G267700 of *Glycine max*; HvAACT1 of *Hordeum vulgare*; MtMATE1, MtMATE2, Medtr1g108840, Medtr1g108990, Medtr1g109060.2, Medtr1g109060.1 of *Medicago truncatula*; NtJAT1, NtMATE1 and NtMATE2 of *Nicotiana tabacum*; SlMTP77 of *Solanum lycopersicum*; SbMATE1 of *Sorghum bicolor*; VvAM1 and VvAM3 of *Vitis vinifera*; VcMATE1 and VcMATE4 of *Vaccinium corymbosum*. The tree was constructed by aligning the protein sequences by Clustal-W and the evolutionary history was inferred using the Neighbour-Joining method. The percentage of replicate trees in which the associated taxa clustered together in the bootstrap test (1000 replicates) are shown next to the branches. The tree is drawn to scale, with branch lengths in the same units as those of the evolutionary distances used to infer the phylogenetic tree. The evolutionary distances were computed using the Poisson correction method and are in the units of the number of amino acid substitutions per site.

At the amino acid level, LaMATE2 exhibits a high similarity to a cluster of MATE proteins, which includes a heterogeneous group of transporters (different substrates, subcellular localization and physiological role). Among the characterized members of this cluster, NtJAT1 and AtDTX1 are the closest homologs of LaMATE2, 55 and 50% of identities, respectively (Figure 3). Both are localized at the plasma membrane (PM) of root cells and function as efflux carriers for plant-derived alkaloids and other toxic compounds (Li et al., 2002; Morita et al., 2009).

Several ABC transporters and MATE proteins mediating the export of compounds into the soil have been described (Weston et al., 2012; Baetz and Martinoia, 2014). However, none of them was shown so far to be responsible for the root release of flavonoids and isoflavonoids. Our data showing a connection between flavonoid exudation and *LaMATE2* expression together with the fact that LaMATE2 fused with GFP co-localizes with the PM marker AtPIP2a-mCherry (Figure 4) prompted us to investigate whether LaMATE2 acts as a flavonoid transporter.

**Figure 4.**
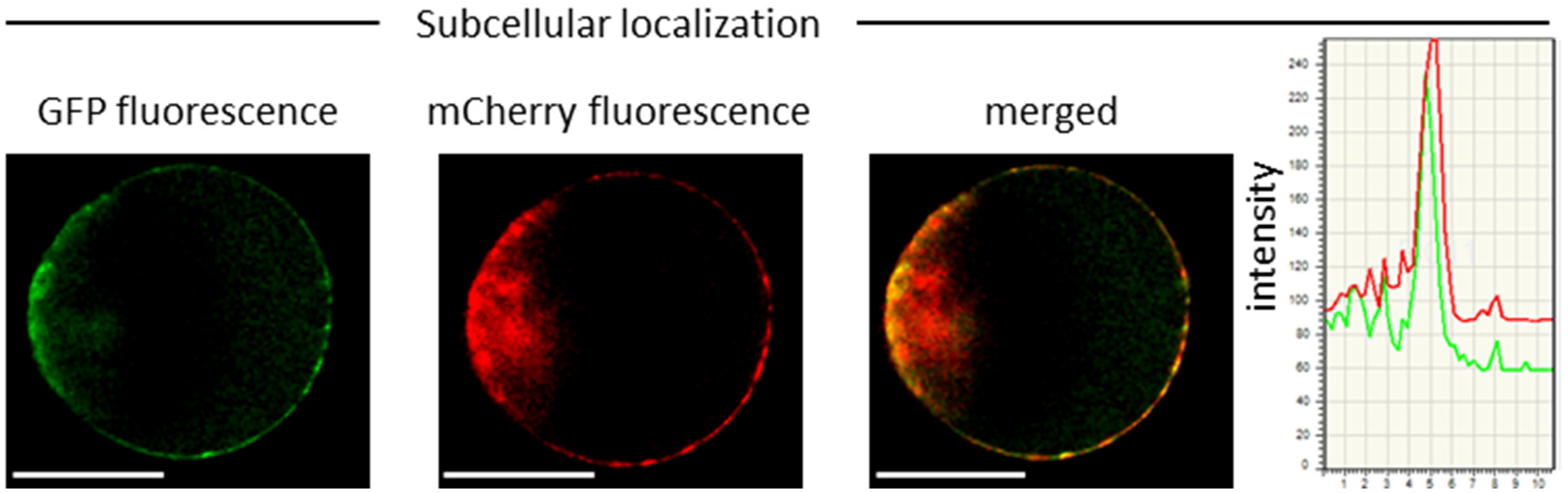
Subcellular localization of LaMATE2. Co-localization of LaMATE2 fused with green fluorescent protein (GFP) and mCherry-labeled plasma membrane marker AtPIP2A in an Arabidopsis mesophyll protoplast. White scale bars = 20 µm. Fluorescence intensity over distance plot of LaMATE2-GFP (green) and AtPIP2a-mCherry (red).

### LaMATE2 silencing reduces genistein release, P mobilization and nodule number

Genistein is one of the major released isoflavonoids of *Fabaceae* inducing *NOD* genes in *Rhizobium* bacteria, attracting them and inducing nodule formation (Zhang and Smith, 1995; Lang et al., 2008; Liu and Murray, 2016). Therefore, to assess whether the LaMATE2 might affect genistein release and nodule formation, we have used RNA-dependent gene silencing (*LaMATE2*-RNAi) in roots of white lupin plants. The effectiveness of the silencing was confirmed by a significant reduction (approximatively −80 and 65 % of *LaMATE2* expression in *LaMATE2*-RNAi transformants grown in N and P deficiency, respectively (Figure 5a, g). Analysis of these roots revealed that the release of genistein was strongly reduced in these transformants (Figure 5b, h) and concomitantly isoflavonoids, the uppermost as glycosides, were accumulating in the cell content of silenced roots, which can be regarded as a direct consequence of the reduced genistein exudation (Figure 5d-f). To investigate whether the impaired genistein exudation influences nodulation, we used the *LaMATE2*-RNAi plants to compare the nodule number of silenced and control plants. Our results highlight the importance of LaMATE2-dependent isoflavonoid release in the early step of nodulation, since *LaMATE2* silencing leads to a highly significant reduction (a. −80 %) of nodules in *LaMATE2*-RNAi compared to empty-vector transformed roots (Figure 5c, 6b, S4). Interestingly exogenous application of genistein onto the LaMATE2-RNAi roots partially restored the nodulation efficiency (Figure 6 a, b). Similarly, *LaMATE2* expression, genistein release patterns and capability of root exudates to mobilize P were measured under P-deficient conditions in silenced plants (Figure 5g-i). Cluster roots of *LaMATE2*-RNAi transformants grown under P deficiency released 85% less genistein along with the lower level of *LaMATE2* expression (a. −60%). The SPE-purified exudates from those roots also exhibited a limited (a. −66%) mobilization capability from a poorly soluble P source (vivianite). These results suggest that LaMATE2 is the major if not the sole genistein exporter from roots of white lupin plants and in P-deficient condition it contributes to the P solubilization process.

**Figure 5.**
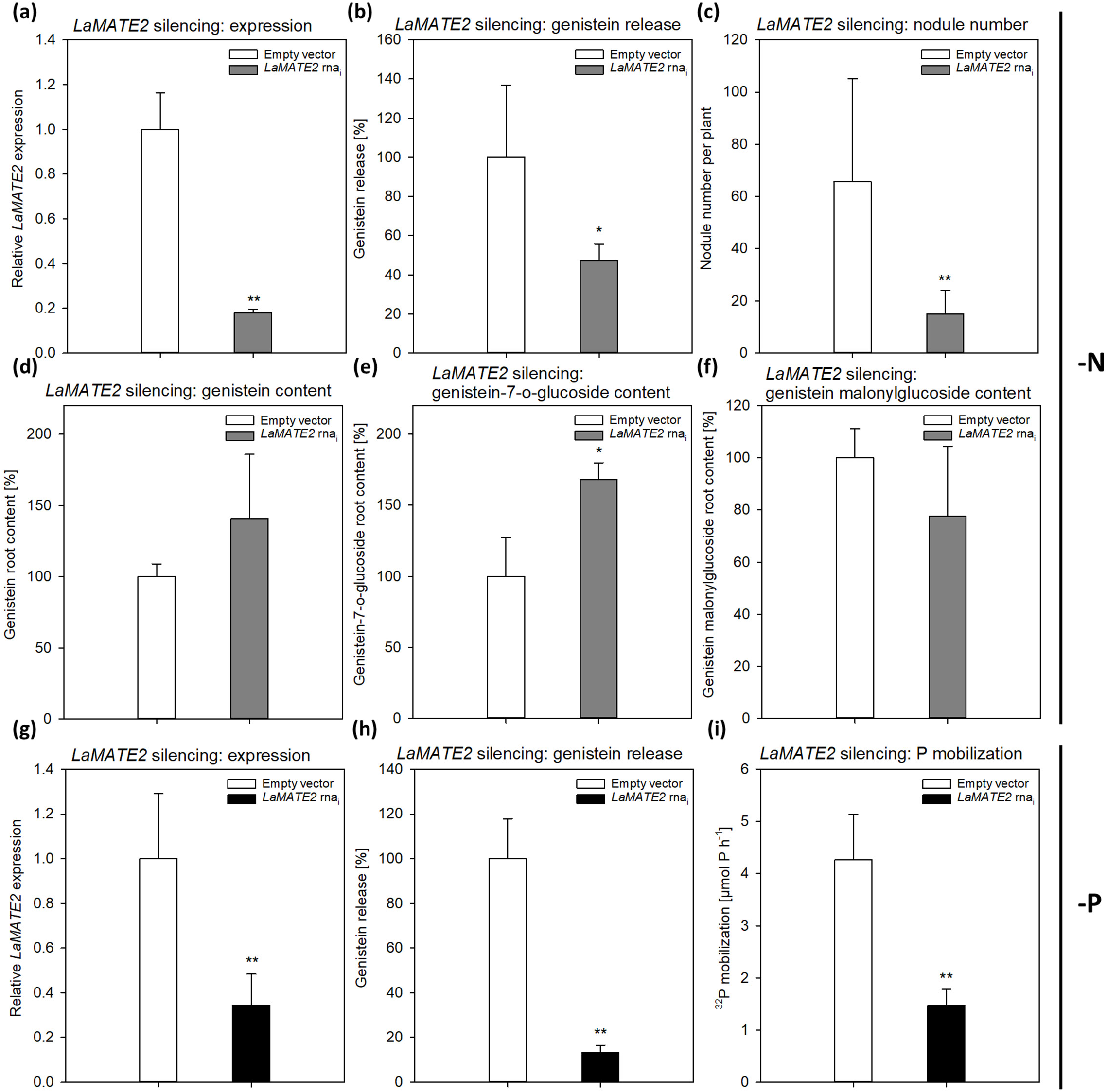
Effect of *LaMATE2* silencing in N and P deficient roots. Alteration of *LaMATE2* expression, flavonoid content and release, nodule number or P mobilization due to *LaMATE2* silencing in N (**a-f**) or P (**g-i**) deficiency. *LaMATE2* relative expression (**a, g**), genistein release (**b, h**), number of nodules per plant (pictures shown in Figure S4) (**c**), cell content: genistein (**d**), genistein 7-O-glucoside (**e**), genistein malonylglucoside (**f**), P mobilization from a poorly soluble P source (**i**). The analyses were performed on roots of N- or P-deficient lupin plants independently transformed with either pRedRoot::*LaMATE2* RNAi (*LaMATE2* RNAi) or empty-vector pRedRoot (Empty vector). All data are expressed relative to Empty-vector-transformed roots, with the exceptions of the number of nodule per plants and P mobilization (µmol P h^−1^). Data are means+SD (* refers to statistically significant differences among the mean value of RNAi and Empty vector, ANOVA Holm–Sidak, N=6-21, *P <0.05; ** for P<0.01).

**Figure 6.**
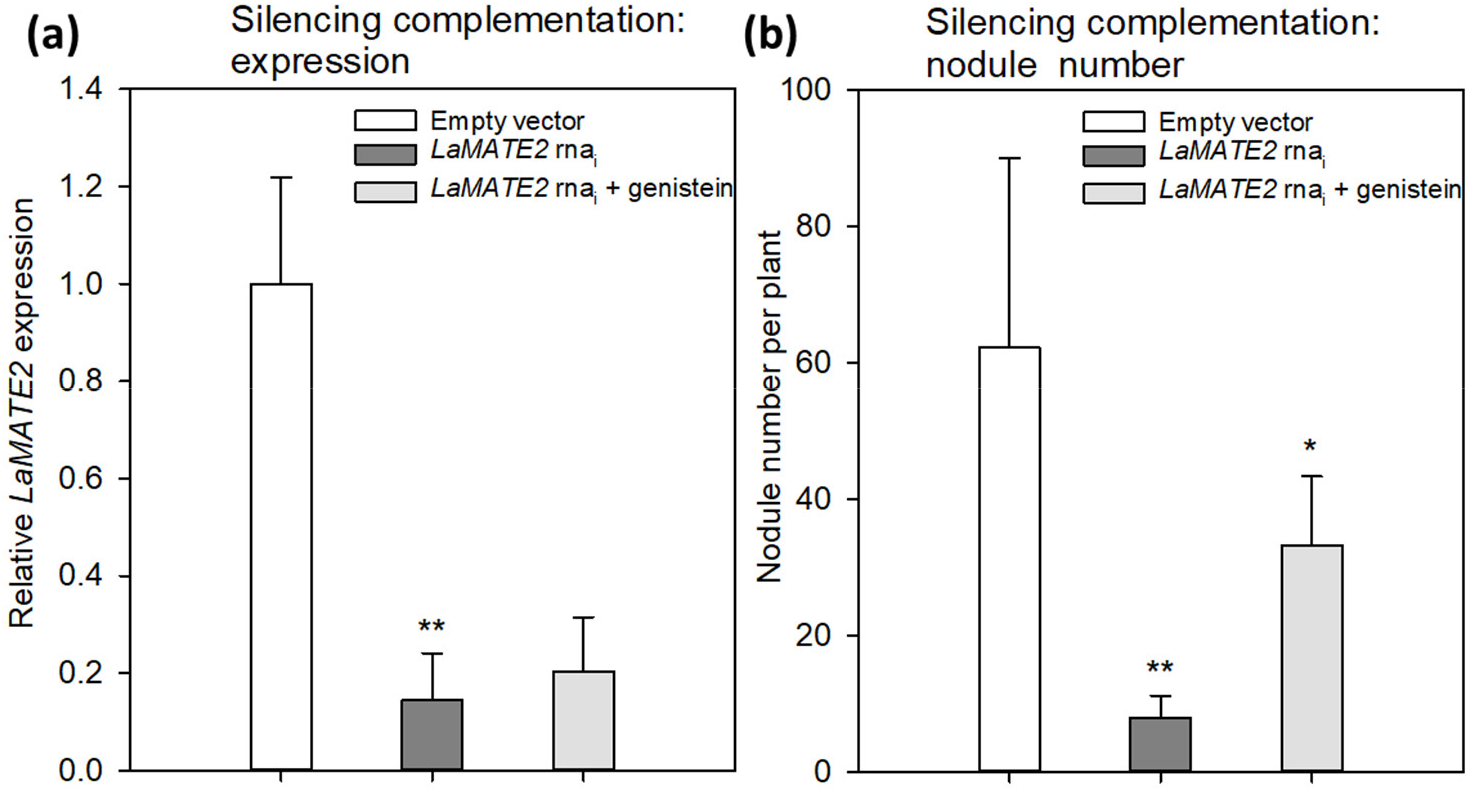
Effect of exogenous genistein on *LaMATE2* expression and nodule numbers. *LaMATE2* relative expression (**a**) and number of nodules per plant (**b**) in roots of N-deficient lupin plants independently transformed with either pRedRoot::*LaMATE2* RNAi (*LaMATE2* RNAi) or empty-vector pRedRoot (Empty vector) or RNAi roots treated with 1 µM genistein (*LaMATE2* RNAi + genistein). Expression data are shown relative to *LaMATE2* expression level in empty vector-transformed roots. Data are means+SD (* refers to statistically significant differences among the mean value of RNAi and EV or RNAi+G values, ANOVA Holm–Sidak, N=6, *P <0.05; ** for P<0.01).

The results showing that genistein export is dependent on the presence of LaMATE2 indicates that within the root this transporter is at least partially localized in the cortex. Attempts in our laboratory to get a more detailed picture on its localization, either using in-situ hybridization or a GUS-promoter construct unfortunately failed. However, it should be mentioned that interestingly, excretion of strigolactones into the soil was independent, whether the transporter was localized specifically in hypodermal passage cells as in Petunia or in the whole cortex as in Medicago (Kretzschmar et al., 2012; Banasiak et al., 2020).

### Transport of phenolics by LaMATE2

To demonstrate that the reduced genistein release of LaMATE2 silenced plants depends directly on the activity of LaMATE2, we expressed it heterologously in *Saccharomyces cerevisiae* and performed transport assays using isolated membrane vesicles (Figure 7). Indeed, using this system, we could observe that LaMATE2 mediates an efficient transport of genistein which was characterized by a saturable kinetic exhibiting an apparent Km of 16.2 µM (Figure 7b). This Km is in the same order of magnitude as determined for the vacuolar flavonoid transporters characterized so far (e.g. AtTT12 and MtMATE1: Km of 50 and 36 µM for epicatechin 3′-O-glucoside, respectively; MtMATE2: Km of 88 µM for cyanidin 3-O-glucoside) (Zhao and Dixon, 2010; Zhao et al., 2011). In plasmalemma vesicles isolated from soybean roots, genistein transport exhibited a Km of 160 µM and was strongly dependent on the presence of ATP (Sugiyama et al., 2007).

**Figure 7.**
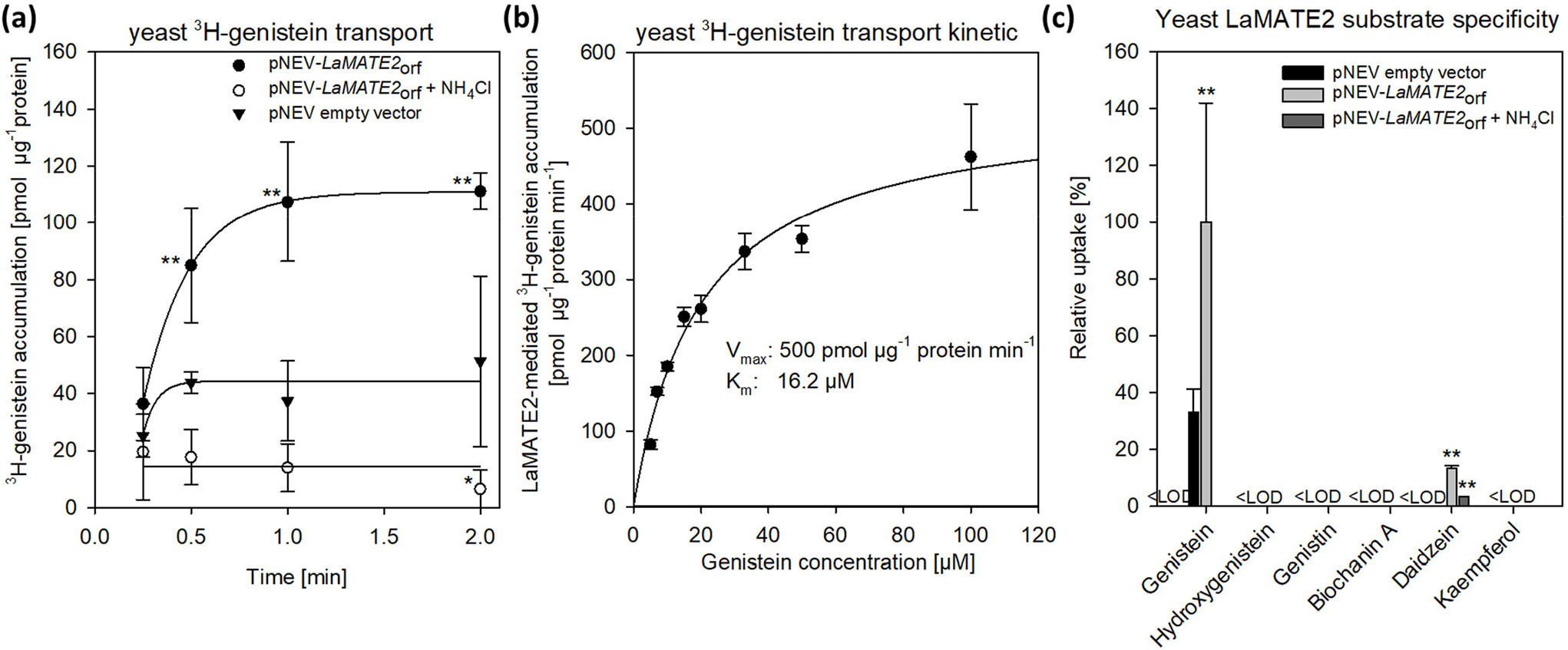
Transport of phenylpropanoid compounds mediated by LaMATE2. LaMATE2-mediated transport rate of phenylpropanoids in yeast microsomal membrane vesicles. (**A**) Time-dependent accumulation of ^3^H-genistein in yeast microsomal membrane vesicles. Membrane vesicles were isolated from yeast transformed with the empty vector (pNEV empty vector) or transformed with pNEV-*LaMATE2*_ORF_. The yeast vesicles were incubated up to 120 s in presence of 5 µM ^3^H-genistein: To determine the effect of the pH gradient, 25 mM NH_4_Cl was added in an assay solution (pNEV-*LaMATE2*_ORF_ + NH_4_Cl), as uncoupler of the proton gradient. (**B**) Concentration-dependent transport of ^3^H-genistein by LaMATE2 in yeast vesicles. Vesicles were incubated for 30 s in the assay solution containing ^3^H-genistein at different concentrations (from 5 to 100 μM). The kinetic parameters of ^3^H-genistein uptake were calculated by subtracting uptake rates recorded in the empty-vector vesicles using the Hanes– Woolf plot. (**C**) Substrate specificity of LaMATE2 transporter was evaluated in presence of genistein, hydroxygenistein, genistin (glycosylated genistein), biochanin A (methylgenistein), daidzein or kaempferol (5 µM, 30 s). All data are expressed relative to pNEV-*LaMATE2*_ORF_-transformed yeast. Data are means±SD of three independent experiments (*refers to statistically significant differences among the mean value in comparison to empty vector sample within each time point; capital letters refer to statistically significant differences among the mean value, ANOVA Holm– Sidak, N=6, *P <0.05, **P <0.01, <LOD: below detectable value).

Up to date, certain MATEs have been shown to exhibit a broad range of substrate specificity (Takanashi et al., 2014), while LaMATE2 displays a strong specificity for the substrate genistein (Figure 6c). Besides genistein, LaMATE2 was able to transport only daidzein at a low rate, but not other tested flavonoids such as genistin (a glycosylated form of genistein), hydroxygenistein, biochanin A (a methylated form of genistein) or kaempferol. Previous observations in soybean showed that the presence of daidzein in the external media strongly limited genistein transport in plasma membrane vesicles of root cells, indicating a possible competition between those molecules (Sugiyama et al., 2007).

A feature of MATE proteins is to couple the substrate transport to an electrochemical gradient, working as H^+^ or Na^+^/substrate antiporters (He et al., 2010; Lu et al., 2013; Tanaka et al., 2013). Our results showed that ATP-dependent ^3^H-genistein accumulation in vesicles deriving from *LaMATE2*_ORF_-expressing yeast was strongly reduced in presence of ammonium chloride, which dissipates the transmembrane proton gradient (Marinova et al., 2007; Zhao and Dixon, 2010) (Figure 7a), indicating that LaMATE2 acts as a substrate-proton co-transporter, similarly to AtTT12 (Marinova et al., 2007), AtDTX1 (Li et al., 2002) and NtJAT1 (Morita et al., 2009).

### Conclusions

In this work the long-sought-after isoflavonoid plasma membrane exporter required to attract symbiotic bacteria for N fixation has been identified. Different *leguminous* plants release different sets of isoflavonoids to induce nodulation (Liu and Murray, 2016). Released genistein has been indicated in faba bean and soybean as *rhizobia* attractant (Zhang and Smith, 1995; Li et al., 2016). Since *Medicago truncatula* and soybean both encode one MATE protein that exhibits high homology to LaMATE2, it is tempting to speculate that these homologues could also act as isoflavonoid exporters, possibly showing slightly different substrate preferences according to the isoflavonoid produced by the plant to initiate the symbiosis. However, it cannot be excluded that in addition to MATE-type transporters and some ABC transporters are also involved in isoflavonoid release. Since isoflavonoids can also release Pi from minerals (Cesco et al., 2010) and organic complexes these transporters may play a dual role in N and P supply.

During the last years, we learned a lot about the signaling pathways and the effectors involved in establishing the legume-*rhizobia* symbiosis. Surprisingly molecular identity of membrane transporters responsible for the release of phenolic compounds initiating interactions was not known. With the identification of LaMATE2 we succeeded to identify the very initial step leading to this symbiosis. Since there is a huge interest to transfer this complex mechanism to other crop plants to grow them in the absence of artificial N-sources we do believe that it is a valuable piece of the puzzle to consider in such an ambitious project.

## ACKNOWLEDGEMENTS

Research was supported by grants from Italian Ministry of University and Research-MIUR (FIRB-Programme Futuro in Ricerca, RBFR08L2ZT and RBFR127WJ9), from funds of the Zurich University; and statutory funds from the Polish Ministry of Science and Higher Education. **Competing interests:** Authors declare no competing interests; and **Data and materials availability:** Sequence data from this article can be found in the EMBL/GenBank data libraries under accession number KY464927, LaMATE2 of *Lupinus albus*.

## AUTHOR CONTRIBUTION

NT, EM, RP, SC, MJ: designed and oversaw the research; WB, LZ, SG, RBF, SV, FV, TM, BB, NT: performed experiments and analyzed data; NT, EM, RP, MJ: wrote the paper

## Supplementary materials

**Figure S1.**
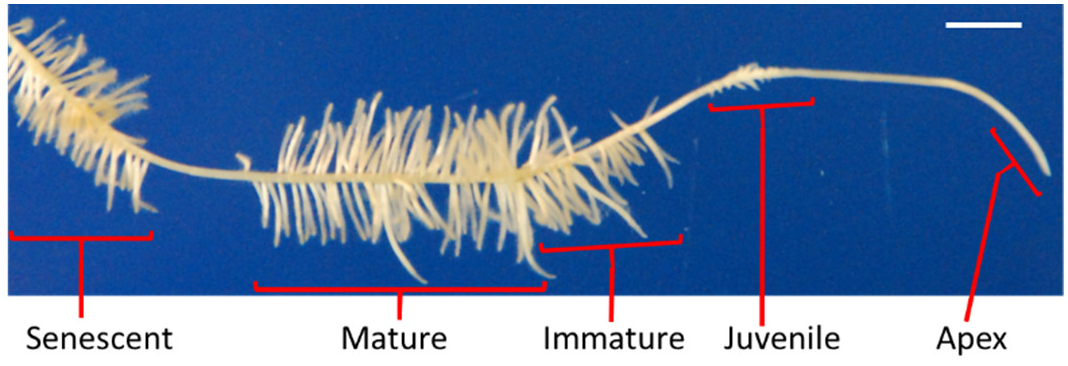
Cluster roots of P-deficient white lupin. Picture of a lateral root of a 4-week-old white lupin grown under P-deficient condition (+N-P). The different developing stages: Juvenile, Immature, Mature, Senescent cluster-root stage and Apex tissue [10 mm from the root tips] are highlighted and named. White scale bar = 10 mm.

**Figure S2.**
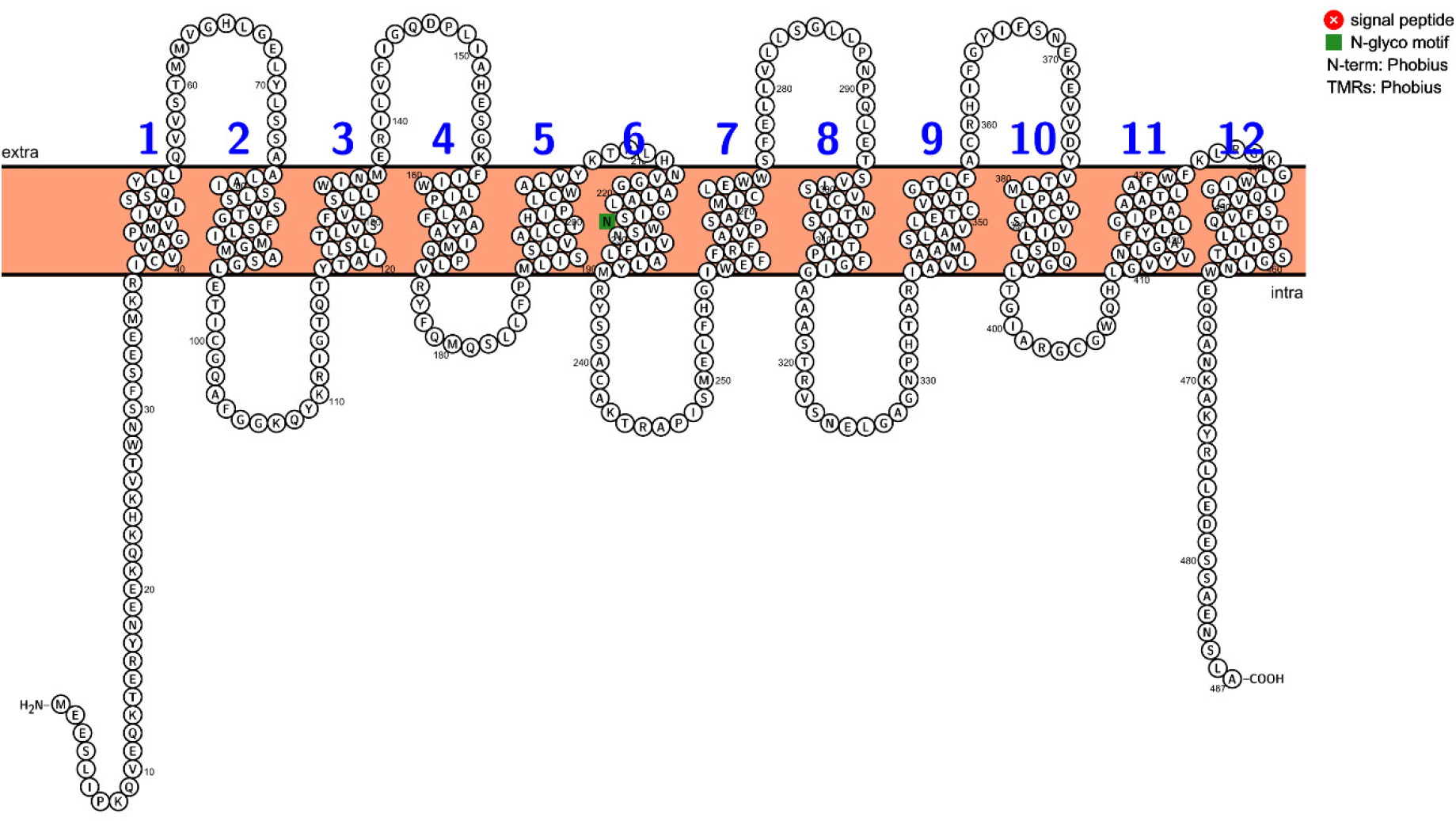
Predicted membrane topology for LaMATE2 from white lupin. The model was obtained by using the PROTTER program (Omasits *et al.,* 2014).

**Figure S3.**
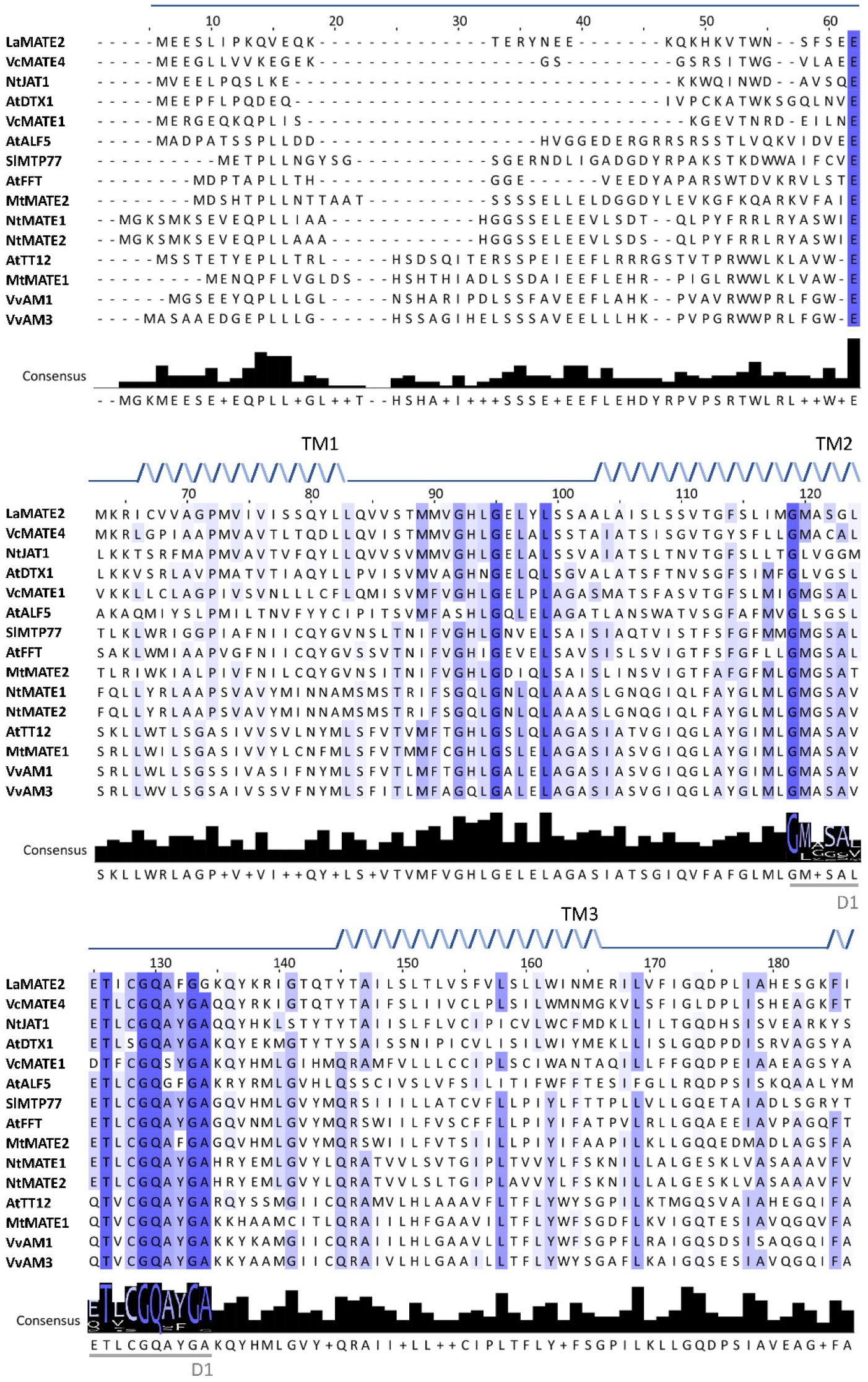
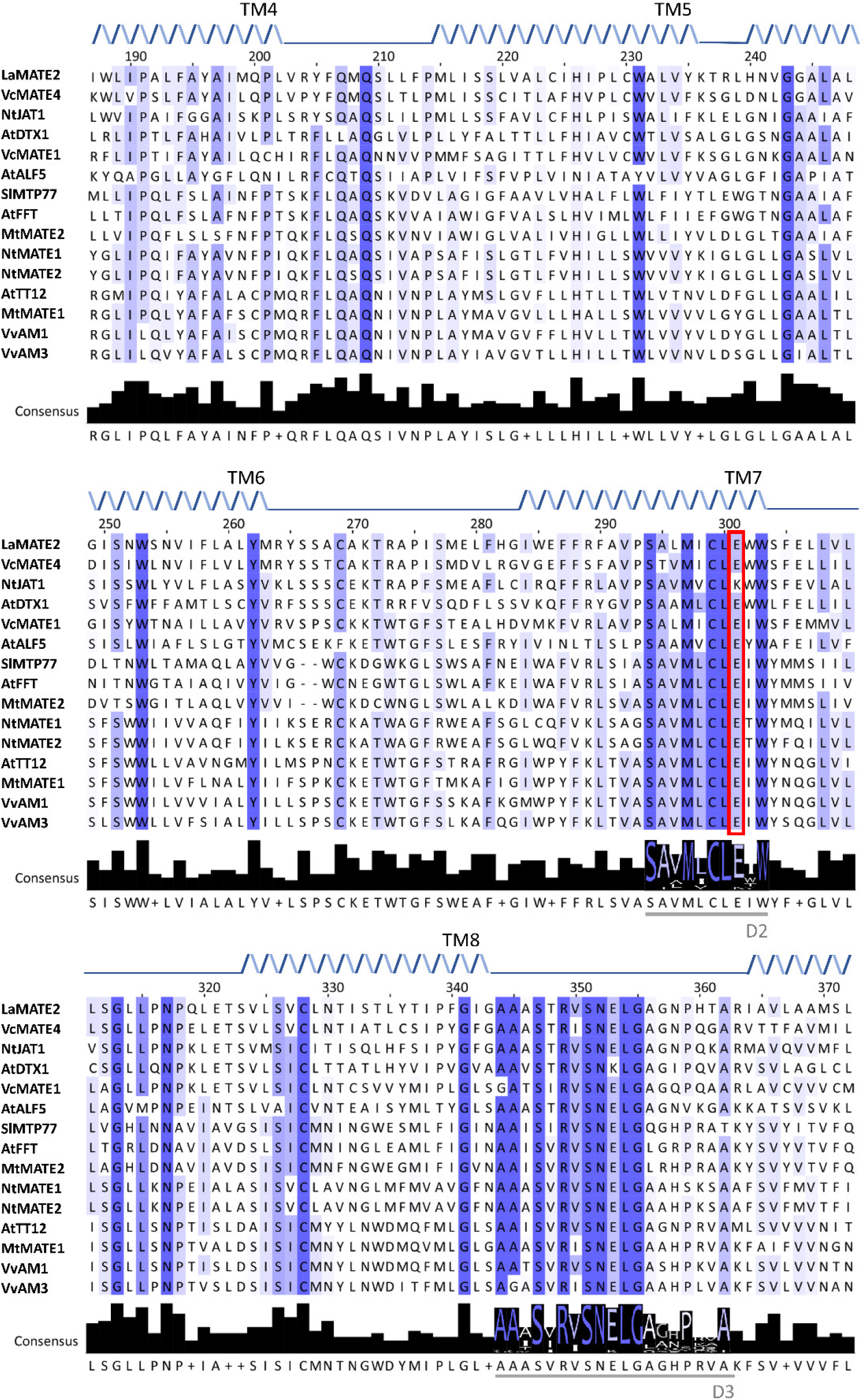
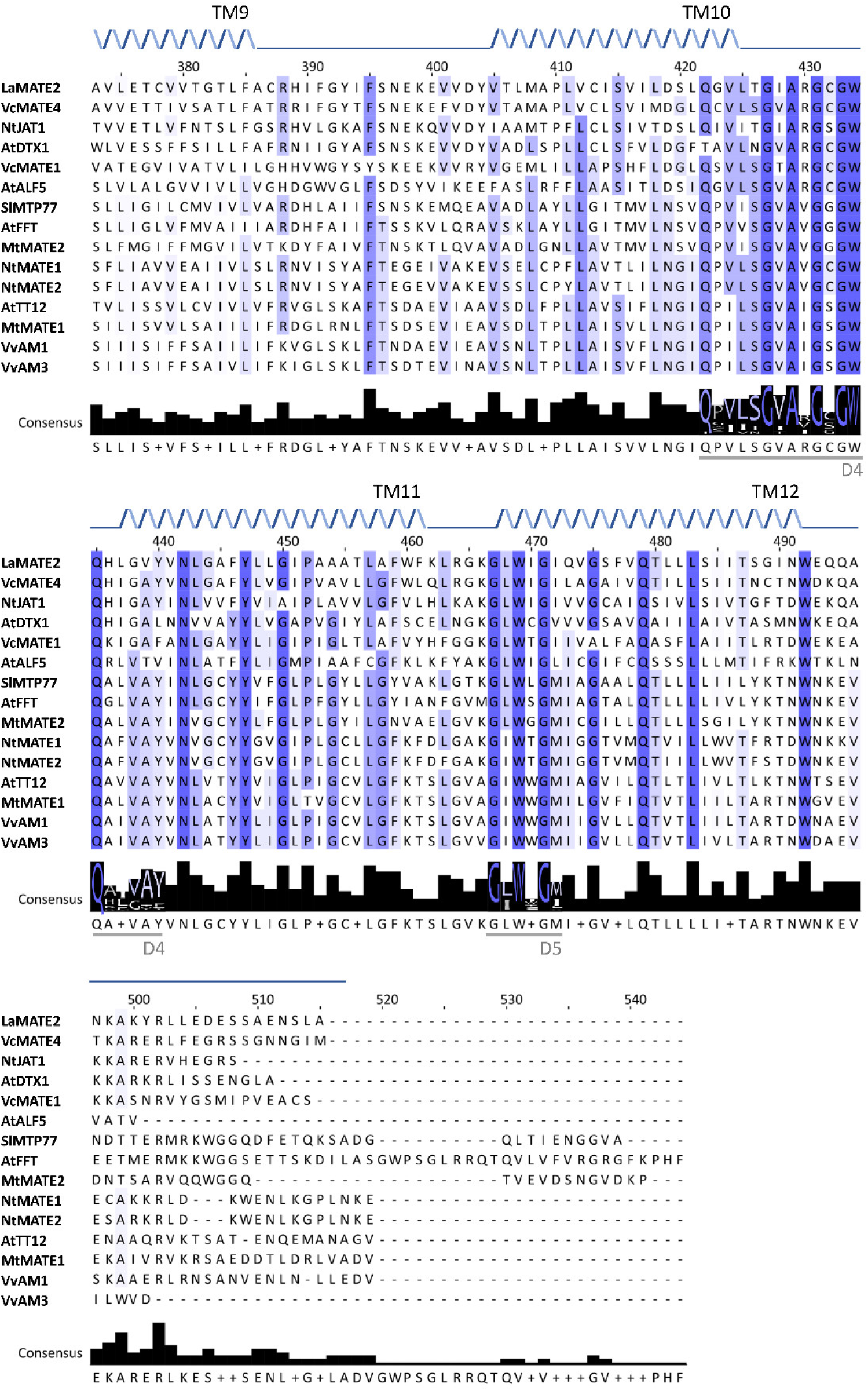
Multiple sequence alignment of amino acid sequence of LaMATE2 with selected MATE transporter orthologs in higher plants. Protein sequence alignment was performed using Clustal-WS using Jalview software version 2 (Waterhouse *et al.,* 2009). Amino acids with only highly conservative substitutions are highlighted in colour blue. Thin grey lines below the consensus sequence (D 1-5) indicate five short stretches of conservative amino acids reported for all 56 Arabidopsis MATE proteins. The red box indicates the residue E290 (TT12), constituting the cation-binding site in the pore. Twelve putative-transmembrane helical domains (TM 1-12) of LaMATE2 are indicated above the alignment

**Figure S4.**
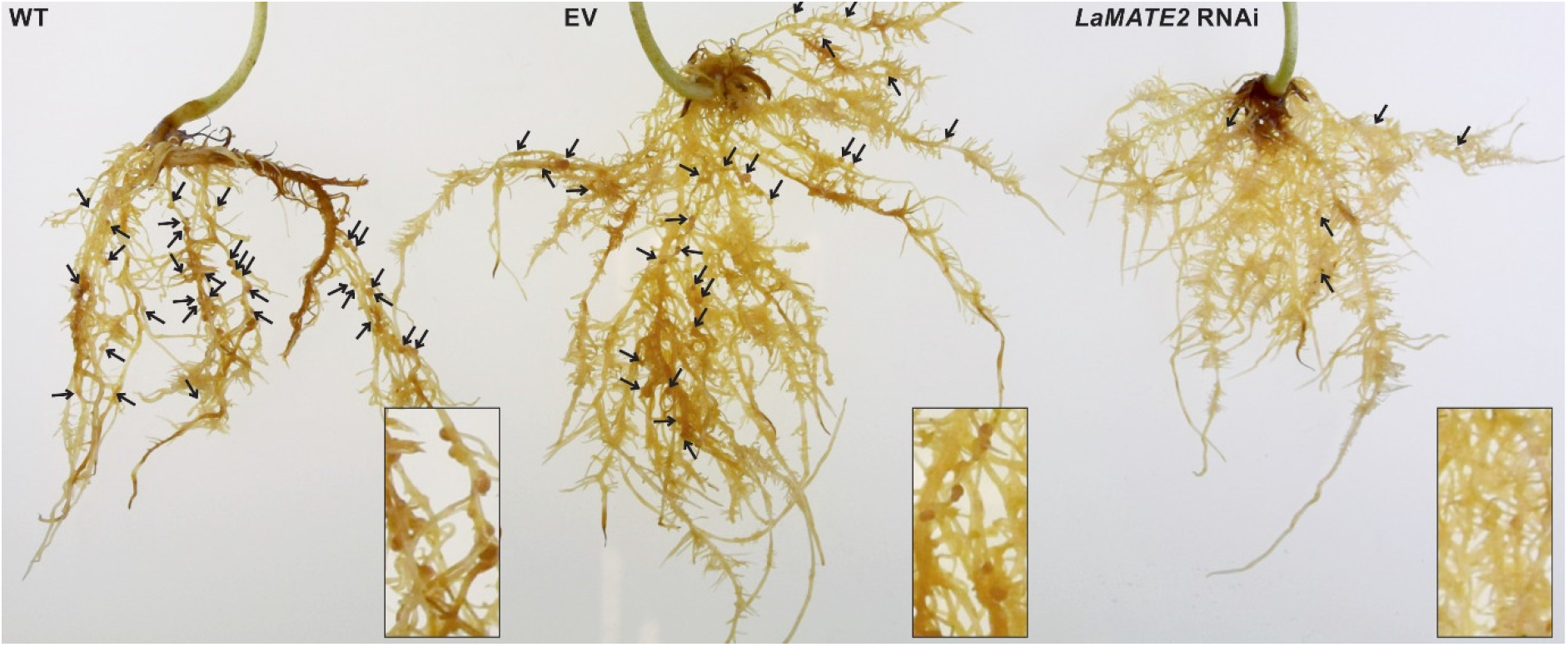
Pictures of *LaMATE2*-silenced roots grown under N-deficient conditions. Roots are shown in the following order: wild type, empty-vector-transformed roots and *LaMATE2*-RNAi-transformed roots. Nodule localizations are highlighted with arrows and close-up inserts are shown in the bottom right corners.

**Figure S5.**
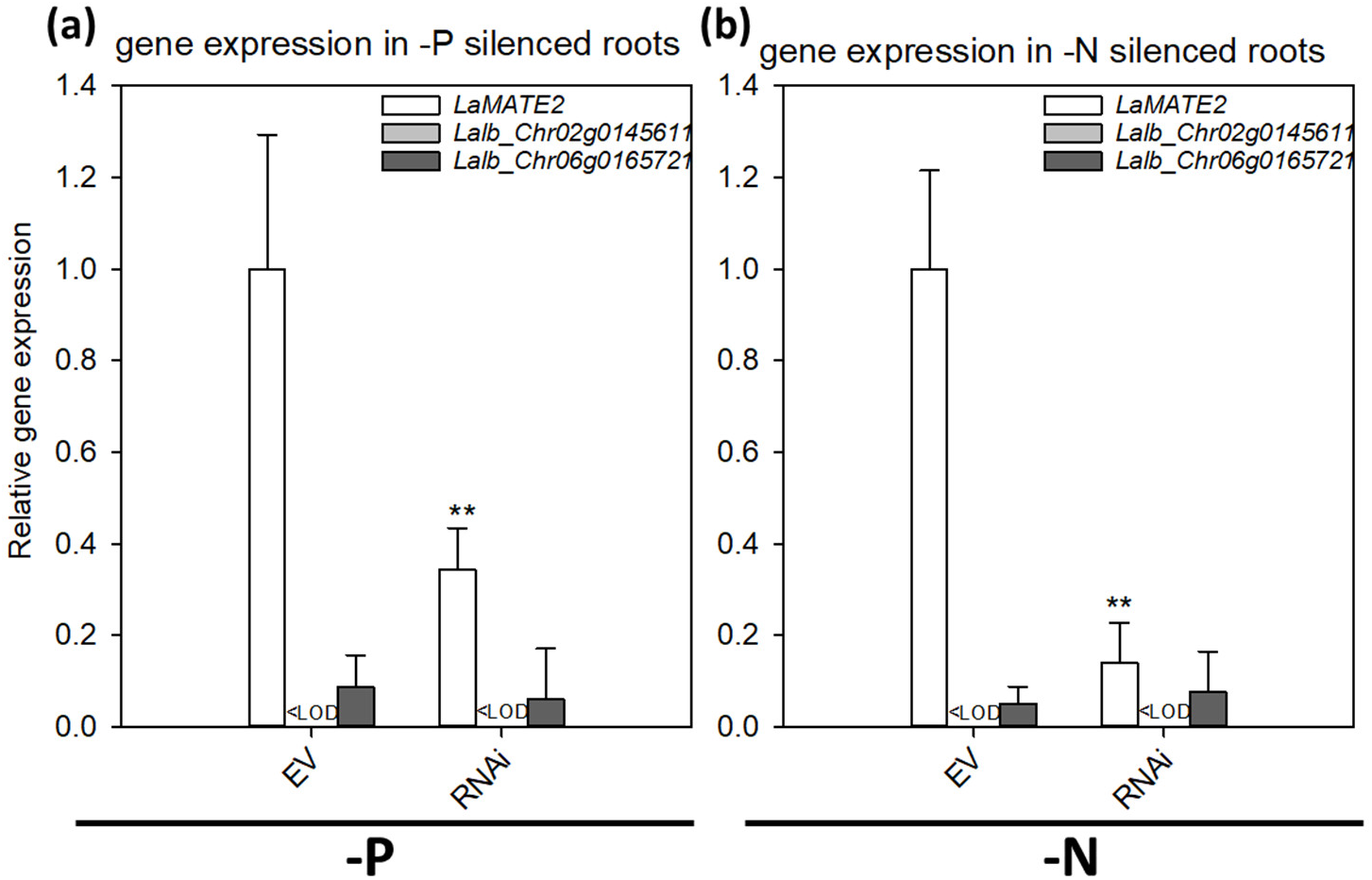
Gene expression in the silencing experiments. Relative expression of *LaMATE2* and the two closest homologues: *Lalb_Chr06g0165721* and *Lalb_Chr02g0145611* in P-deficient (**a**) and N-deficient (**b**) condition in pRedRoot::*LaMATE2* RNAi (RNAi) or empty-vector pRedRoot (EV) transformed roots. Expression data are expressed relative to *LaMATE2* expression in EV-transformed roots. Data are means+SD (* refers to statistically significant differences among the mean value of RNAi and EV, ANOVA Holm–Sidak, N=3, **P <0.01).

**Table S1.**
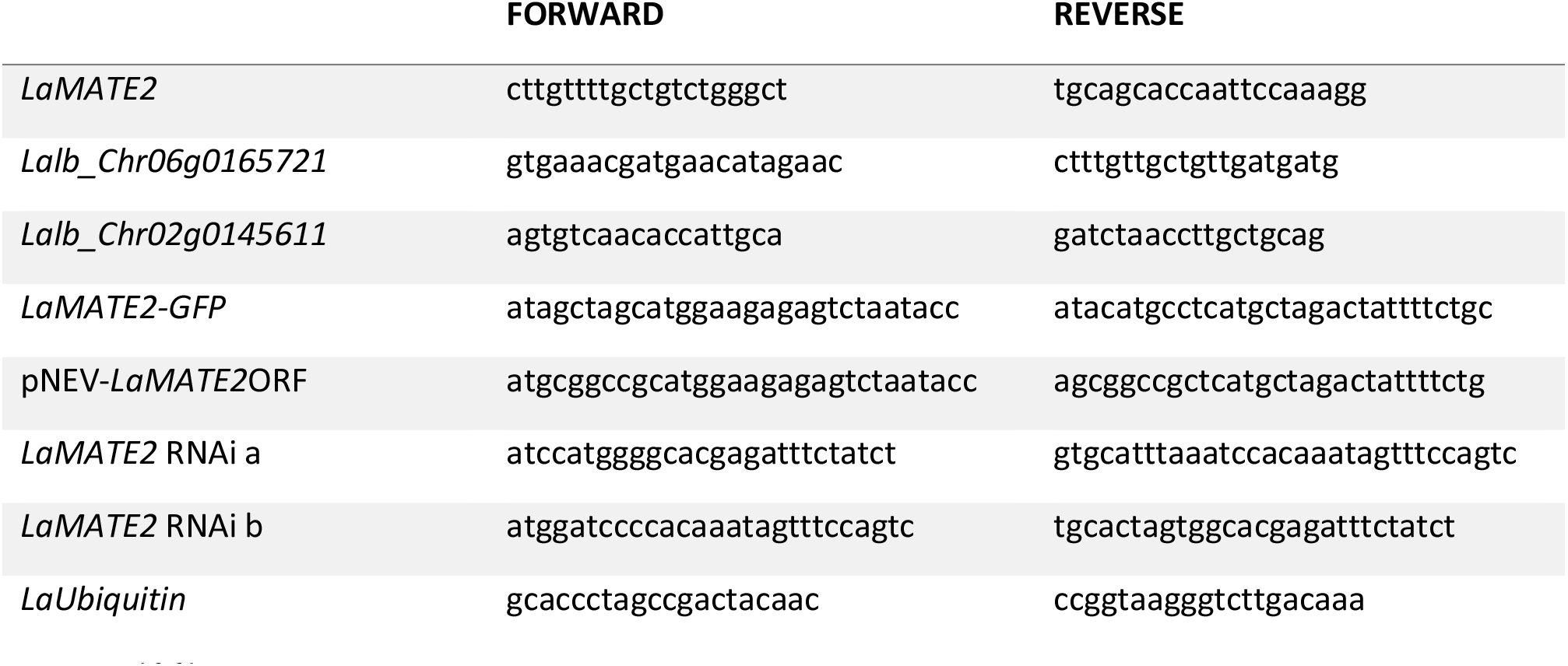
List of primers

## REFERENCES

Baetz, U., and Martinoia, E. (2014). Root exudates: the hidden part of plant defense. Trends in Plant Science 19, 90–98.

Banasiak, J., Borghi, L., Stec, N., Martinoia, E., and Jasiński, M. (2020). The Full-Size ABCG Transporter of *Medicago truncatula* Is Involved in Strigolactone Secretion, Affecting Arbuscular Mycorrhiza. 11.

Biala, W., Banasiak, J., Jarzyniak, K., Pawela, A., and Jasinski, M. (2017). Medicago truncatula ABCG10 is a transporter of 4-coumarate and liquiritigenin in the medicarpin biosynthetic pathway. Journal of experimental botany 68, 3231–3241.

Burke, D., Dawson, D., and Stearns, T. (2000). Methods in yeast genetics: a Cold Spring Harbor Laboratory course manual, 2000 ed. Cold Spring Harbor Press: Plainview, NY 17, 205.

Cesco, S., Neumann, G., Tomasi, N., Pinton, R., and Weisskopf, L. (2010). Release of plant-borne flavonoids into the rhizosphere and their role in plant nutrition. Plant and Soil 329, 1–25.

Durrett, T.P., Gassmann, W., and Rogers, E.E. (2007). The FRD3-Mediated Efflux of Citrate into the Root Vasculature Is Necessary for Efficient Iron Translocation. Plant Physiology 144, 197–205.

Furukawa, J., Yamaji, N., Wang, H., Mitani, N., Murata, Y., Sato, K., Katsuhara, M., Takeda, K., and Ma, J.F. (2007). An Aluminum-Activated Citrate Transporter in Barley. Plant and Cell Physiology 48, 1081–1091.

Gietz, R.D., and Woods, R.A. (2002). Transformation of yeast by lithium acetate/single-stranded carrier DNA/polyethylene glycol method. Methods in enzymology 350, 87–96.

He, X., Szewczyk, P., Karyakin, A., Evin, M., Hong, W.-X., Zhang, Q., and Chang, G. (2010). Structure of a cation-bound multidrug and toxic compound extrusion transporter. Nature 467, 991.

Hufnagel, B., Marques, A., Soriano, A., Marquès, L., Divol, F., Doumas, P., Sallet, E., Mancinotti, D., Carrere, S., Marande, W., Arribat, S., Keller, J., Huneau, C., Blein, T., Aimé, D., Laguerre, M., Taylor, J., Schubert, V., Nelson, M., Geu-Flores, F., Crespi, M., Gallardo, K., Delaux, P.-M., Salse, J., Bergès, H., Guyot, R., Gouzy, J., and Péret, B. (2020). High-quality genome sequence of white lupin provides insight into soil exploration and seed quality. Nature Communications 11, 492.

Jiang, Y., Wang, W., Xie, Q., Liu, N., Liu, L., Wang, D., Zhang, X., Yang, C., Chen, X., Tang, D., and Wang, E. (2017). Plants transfer lipids to sustain colonization by mutualistic mycorrhizal and parasitic fungi. Science 356, 1172–1175.

Jin, J.B., Kim, Y.A., Kim, S.J., Lee, S.H., Kim, D.H., Cheong, G.W., and Hwang, I. (2001). A new dynamin-like protein, ADL6, is involved in trafficking from the *trans*-Golgi network to the central vacuole in Arabidopsis. The Plant cell 13, 1511–1526.

Klein, M., Mamnun, Y.M., Eggmann, T., Schüller, C., Wolfger, H., Martinoia, E., and Kuchler, K. (2002). The ATP-binding cassette (ABC) transporter Bpt1p mediates vacuolar sequestration of glutathione conjugates in yeast. FEBS letters 520, 63–67.

Komarova, N.Y., Thor, K., Gubler, A., Meier, S., Dietrich, D., Weichert, A., Suter Grotemeyer, M., Tegeder, M., and Rentsch, D. (2008). AtPTR1 and AtPTR5 Transport Dipeptides in Planta. Plant Physiology 148, 856–869.

Koressaar, T., and Remm, M. (2007). Enhancements and modifications of primer design program Primer3. Bioinformatics 23, 1289–1291.

Kretzschmar, T., Kohlen, W., Sasse, J., Borghi, L., Schlegel, M., Bachelier, J.B., Reinhardt, D., Bours, R., Bouwmeester, H.J., and Martinoia, E. (2012). A petunia ABC protein controls strigolactone-dependent symbiotic signalling and branching. Nature 483, 341–344.

Lambers, H., Martinoia, E., and Renton, M. (2015). Plant adaptations to severely phosphorus-impoverished soils. Current Opinion in Plant Biology 25, 23–31.

Lang, K., Lindemann, A., Hauser, F., and Göttfert, M. (2008). The genistein stimulon of *Bradyrhizobium japonicum*. Molecular Genetics and Genomics 279, 203–211.

Li, B., Li, Y.-Y., Wu, H.-M., Zhang, F.-F., Li, C.-J., Li, X.-X., Lambers, H., and Li, L. (2016). Root exudates drive interspecific facilitation by enhancing nodulation and N_2_ fixation. Proceedings of the National Academy of Sciences 113, 6496–6501.

Li, L., He, Z., Pandey, G.K., Tsuchiya, T., and Luan, S. (2002). Functional Cloning and Characterization of a Plant Efflux Carrier for Multidrug and Heavy Metal Detoxification. Journal of Biological Chemistry 277, 5360–5368.

Limpens, E., Ramos, J., Franken, C., Raz, V., Compaan, B., Franssen, H., Bisseling, T., and Geurts, R. (2004). RNA interference in *Agrobacterium rhizogenes*-transformed roots of *Arabidopsis* and *Medicago truncatula*. Journal of Experimental Botany 55, 983–992.

Liu, C.-W., and Murray, J. (2016). The Role of Flavonoids in Nodulation Host-Range Specificity: An Update. Plants 5, 33.

Livak, K.J., and Schmittgen, T.D. (2001). Analysis of Relative Gene Expression Data Using Real-Time Quantitative PCR and the 2−ΔΔCT Method. Methods 25, 402–408.

Lu, M., Radchenko, M., Symersky, J., Nie, R., and Guo, Y. (2013). Structural insights into H^+^-coupled multidrug extrusion by a MATE transporter. Nature Structural & Molecular Biology 20, 1310–1317.

Luginbuehl, L.H., Menard, G.N., Kurup, S., Van Erp, H., Radhakrishnan, G.V., Breakspear, A., Oldroyd, G.E.D., and Eastmond, P.J. (2017). Fatty acids in arbuscular mycorrhizal fungi are synthesized by the host plant. Science 356, 1175–1178.

Magalhaes, J.V., Liu, J., Guimaraes, C.T., Lana, U.G.P., Alves, V.M.C., Wang, Y.H., Schaffert, R.E., Hoekenga, O.A., Pineros, M.A., Shaff, J.E., Klein, P.E., Carneiro, N.P., Coelho, C.M., Trick, H.N., and Kochian, L.V. (2007). A gene in the multidrug and toxic compound extrusion (MATE) family confers aluminum tolerance in sorghum. Nature Genetics 39, 1156–1161.

Marinova, K., Kleinschmidt, K., Weissenbock, G., and Klein, M. (2007). Flavonoid Biosynthesis in Barley Primary Leaves Requires the Presence of the Vacuole and Controls the Activity of Vacuolar Flavonoid Transport. Plant Physiology 144, 432–444.

Martin, F.M., Uroz, S., and Barker, D.G. (2017). Ancestral alliances: Plant mutualistic symbioses with fungi and bacteria. Science 356, eaad4501.

Massonneau, A., Langlade, N., Leon, S., Smutny, J., Vogt, E., Neumann, G., and Martinoia, E. (2001). Metabolic changes associated with cluster root development in white lupin (*Lupinus albus* L.): relationship between organic acid excretion, sucrose metabolism and energy status. Planta 213, 534–542.

Morita, M., Shitan, N., Sawada, K., Van Montagu, M.C.E., Inzé, D., Rischer, H., Goossens, A., Oksman-Caldentey, K.-M., Moriyama, Y., and Yazaki, K. (2009). Vacuolar transport of nicotine is mediated by a multidrug and toxic compound extrusion (MATE) transporter in *Nicotiana tabacum*. Proceedings of the National Academy of Sciences of the United States of America 106, 2447–2452.

Nelson, B.K., Cai, X., and Nebenführ, A. (2007). A multicolored set of in vivo organelle markers for co-localization studies in *Arabidopsis* and other plants. The Plant Journal 51, 1126–1136.

Neumann, G., and Martinoia, E. (2002). Cluster roots - an underground adaptation for survival in extreme environments. Trends in Plant Science 7, 162–167.

Omasits, U., Ahrens, C.H., Müller, S., and Wollscheid, B. (2014). Protter: interactive protein feature visualization and integration with experimental proteomic data. Bioinformatics 30, 884–886.

Purnell, H.M. (1960). Studies of the family Proteaceae. Anatomy and morphology of the roots of some Victorian species. Australian Journal of Botany 8, 38–50.

Quandt, H.J., Pühler, A., and Broer, I. (1993). Transgenic root nodules of *Vicia hirsuta:* a fast and efficient system for the study of gene expression in indeterminate-type nodules. MPMI-Molecular Plant Microbe Interactions 6, 699–706.

Ritz, C., and Spiess, A.N. (2008). qpcR: an R package for sigmoidal model selection in quantitative real-time polymerase chain reaction analysis. Bioinformatics 24, 1549–1551.

Sauer, N., and Stolz, J. (1994). SUC1 and SUC2: two sucrose transporters from *Arabidopsis thaliana*; expression and characterization in baker’s yeast and identification of the histidine-tagged protein. The Plant Journal 6, 67–77.

Staszków, A., Swarcewicz, B., Banasiak, J., Muth, D., Jasiński, M., and Stobiecki, M. (2011). LC/MS profiling of flavonoid glycoconjugates isolated from hairy roots, suspension root cell cultures and seedling roots of Medicago truncatula. Metabolomics 7, 604–613.

Stępkowski, T., Żak, M., Moulin, L., Króliczak, J., Golińska, B., Narożna, D., Safronova, V.I., and Mądrzak, C.J. (2011). *Bradyrhizobium canariense* and *Bradyrhizobium japonicum* are the two dominant *rhizobium* species in root nodules of lupin and serradella plants growing in Europe. Systematic and Applied Microbiology 34, 368–375.

Strozycki, P.M., Skąpska, A., Szcześniak, K., Sobieszczuk, E., Briat, J.-F., and Legocki, A.B. (2003). Differential expression and evolutionary analysis of the three ferritin genes in the legume plant *Lupinus luteus*. Physiologia Plantarum 118, 380–389.

Subramanian, S., Stacey, G., and Yu, O. (2006). Endogenous isoflavones are essential for the establishment of symbiosis between soybean and *Bradyrhizobium japonicum*. The Plant Journal 48, 261–273.

Sugiyama, A., Shitan, N., and Yazaki, K. (2007). Involvement of a soybean ATP-binding cassette-Type transporter in the secretion of genistein, a signal flavonoid in legume-*Rhizobium* Symbiosis. Plant Physiology 144, 2000–2008.

Takanashi, K., Shitan, N., and Yazaki, K. (2014). The multidrug and toxic compound extrusion (MATE) family in plants. Plant Biotechnology 31, 417–430.

Tamura, K., Stecher, G., Peterson, D., Filipski, A., and Kumar, S. (2013). MEGA6: Molecular Evolutionary Genetics Analysis version 6.0. Molecular biology and evolution 30, 2725–2729.

Tanaka, Y., Hipolito, C.J., Maturana, A.D., Ito, K., Kuroda, T., Higuchi, T., Katoh, T., Kato, H.E., Hattori, M., Kumazaki, K., Tsukazaki, T., Ishitani, R., Suga, H., and Nureki, O. (2013). Structural basis for the drug extrusion mechanism by a MATE multidrug transporter. Nature 496, 247–251.

Tilman, D. (1999). Global environmental impacts of agricultural expansion: The need for sustainable and efficient practices. Proceedings of the National Academy of Sciences 96, 5995–6000.

Tilman, D., Fargione, J., Wolff, B., D’antonio, C., Dobson, A., Howarth, R., Schindler, D., Schlesinger, W.H., Simberloff, D., and Swackhamer, D. (2001). Forecasting Agriculturally Driven Global Environmental Change. Science 292, 281–284.

Udvardi, M., and Poole, P.S. (2013). Transport and Metabolism in Legume-*Rhizobia* Symbioses. Annual Review of Plant Biology 64, 781–805.

Uhde-Stone, C., Liu, J., Zinn, K.E., Allan, D.L., and Vance, C.P. (2005). Transgenic proteoid roots of white lupin: a vehicle for characterizing and silencing root genes involved in adaptation to P stress. Plant Journal 44, 840–853.

Untergasser, A., Cutcutache, I., Koressaar, T., Ye, J., Faircloth, B.C., Remm, M., and Rozen, S.G. (2012). Primer3—new capabilities and interfaces. Nucleic Acids Research 40, e115.

Vance, C.P., Uhde-Stone, C., and Allan, D.L. (2003). Phosphorus acquisition and use: critical adaptations by plants for securing a nonrenewable resource. New Phytologist 157, 423–447.

Wang, Z., Straub, D., Yang, H., Kania, A., Shen, J., Ludewig, U., and Neumann, G. (2014). The regulatory network of cluster-root function and development in phosphate-deficient white lupin (*Lupinus albus*) identified by transcriptome sequencing. Physiologia Plantarum 151, 323–338.

Wasson, A.P., Pellerone, F.I., and Mathesius, U. (2006). Silencing the flavonoid pathway in *Medicago truncatula* inhibits root nodule formation and prevents auxin transport regulation by rhizobia. Plant Cell 18, 1617–1629.

Waterhouse, A.M., Procter, J.B., Martin, D.M.A., Clamp, M., and Barton, G.J. (2009). Jalview Version 2—a multiple sequence alignment editor and analysis workbench. Bioinformatics 25, 1189–1191.

Weisskopf, L., Tomasi, N., Santelia, D., Martinoia, E., Langlade, N.B., Tabacchi, R., and Abou-Mansour, E. (2006). Isoflavonoid exudation from white lupin roots is influenced by phosphate supply, root type and cluster-root stage. New Phytologist 171, 657–668.

Weston, L.A., Ryan, P.R., and Watt, M. (2012). Mechanisms for cellular transport and release of allelochemicals from plant roots into the rhizosphere. Journal of Experimental Botany.

Zhang, F., and Smith, D.L. (1995). Preincubation of Bradyrhizobium japonicum with Genistein Accelerates Nodule Development of Soybean at Suboptimal Root Zone Temperatures. Plant Physiology 108, 961–968.

Zhao, J. (2015). Flavonoid transport mechanisms: how to go, and with whom. Trends in Plant Science 20, 576–585.

Zhao, J., and Dixon, R.A. (2010). The ‘ins’ and ‘outs’ of flavonoid transport. Trends in Plant Science 15, 72–80.

Zhao, J., Huhman, D., Shadle, G., He, X.-Z., Sumner, L.W., Tang, Y., and Dixon, R.A. (2011). MATE2 Mediates Vacuolar Sequestration of Flavonoid Glycosides and Glycoside Malonates in *Medicago truncatula*. The Plant Cell 23, 1536–1555.

